# *E. coli* FtsN coordinates synthesis and degradation of septal peptidoglycan by partitioning between a synthesis track and a denuded glycan track

**DOI:** 10.1101/2024.05.13.594014

**Authors:** Zhixin Lyu, Xinxing Yang, Atsushi Yahashiri, Stephen Ha, Joshua W. McCausland, Xinlei Chen, Brooke M. Britton, David S. Weiss, Jie Xiao

## Abstract

The *E. coli* cell division protein FtsN was proposed to coordinate septal peptidoglycan (sPG) synthesis and degradation to ensure robust cell wall constriction without lethal lesions. Although the precise mechanism remains unclear, previous work highlights the importance of two FtsN domains: the E domain, which interacts with and activates the sPG synthesis complex FtsWIQLB, and the SPOR domain, which binds to denuded glycan (dnG) strands, key intermediates in sPG degradation. Here, we used single-molecule tracking of FtsN and FtsW (a proxy for the sPG synthesis complex FtsWIQLB) to investigate how FtsN coordinates the two opposing processes. We observed dynamic behaviors indicating that FtsN’s SPOR domain binds to dnGs cooperatively, which both sequesters the sPG synthesis complex on dnG (termed as the dnG-track) and protects dnGs from degradation by lytic transglycosylases (LTs). The release of the SPOR domain from dnGs leads to activating the sPG synthesis complex on the sPG-track and simultaneously exposing those same dnGs to degradation. Furthermore, FtsN’s SPOR domain self-interacts and facilitates the formation of a multimeric sPG synthesis complex on both tracks. The cooperative self-interaction of the SPOR domain creates a sensitive switch to regulate the partitioning of FtsN between the dnG- and sPG-tracks, thereby controlling the balance between sequestered and active populations of the sPG synthesis complex. As such, FtsN coordinates sPG synthesis and degradation in space and time.

## Introduction

Gram-negative (diderm) bacteria such as *Escherichia coli* have a rigid cell wall composed of a thin peptidoglycan (PG) layer. The PG is made of repeated N-acetylglucosamine (GlcNAc)-N-acetylmuramic acid (MurNAc) units cross-linked via short peptide bridges^1^. This thin PG layer is constantly remodeled by a large collection of cell wall synthases and hydrolases to allow cell growth and division. In *E. coli*, constriction requires an essential septal PG (sPG) synthase complex formed by the glycosyltransferase FtsW and the transpeptidase FtsI, along with more than ten nonessential PG hydrolases, including three periplasmic amidases (Ami) and eight known and putative lytic transglycosylases (LTs)^2, 3^. Amidases remove peptide crosslinks by cleaving the peptide moiety from MurNAc, which generates denuded glycan (dnG) strands^4, 5^. dnGs are subsequently destroyed by LTs, which cleave the β-1,4-glycosidic bond between MurNAc and GlcNAc^2, 6, 7^. The degradation of PG must be precisely regulated and synchronized with new sPG synthesis in both space and time to ensure robust cell wall constriction without causing lethal lesions. However, the mechanisms underlying this spatiotemporal coordination remain unclear.

FtsN is an essential bitopic membrane protein in *E. coli* and other γ-proteobacteria. It possesses four functional domains with distinct roles in cell division: (1) a short, N-terminal cytoplasmic domain (FtsN^Cyto^) that directly interacts with the early division protein FtsA to trigger the activation of FtsW^8–10^; (2) an α-helical transmembrane domain (FtsN^TM^) that anchors FtsN to the inner membrane^11^; (3) a short, loosely-formed helical essential domain (FtsN^E^) in the periplasm that binds to FtsI and its scaffolding protein FtsL to activate the sPG synthesis complex formed by FtsWI and FtsQLB^12–15^; and (4) a periplasmic, highly conserved C-terminal SPOR domain (FtsN^SPOR^) that binds to dnG^12, 14, 16–19^. A notable feature of FtsN is its long, largely disordered linker between FtsN^TM^ and FtsN^SPOR^, which also includes FtsN^E^. When fully extended, this linker can reach ∼70 nm in length^14^.

In previous single-molecule tracking (SMT) studies in *E. coli*, we observed that FtsW, FtsI, and FtsB (and hence most likely the full sPG synthesis complex formed by FtsWIQLB) exhibit two distinct directional moving populations. One population moves fast at ∼30 nm s^-1^, is inactive in sPG synthesis, and is driven by FtsZ’s treadmilling dynamics through a Brownian ratchet mechanism^15, 20–22^. We term this fast-moving population as on the Z-track. The other population moves slowly at ∼8-9 nm s^-1^, is driven by active sPG synthesis, and is independent of FtsZ’s treadmilling dynamics^15, 20, 21^. We term this slow-moving population as on the sPG-track. Additionally, we observed stationary populations of FtsW, FtsI and FtsB, which we identified as either trapped in the middle of FtsZ polymers or “poised” for synthesis. Thus, SMT provides a powerful means to link the moving dynamics of cell wall synthesis enzymes and regulators to their activities and functions in live bacterial cells.

We found that modulating a cell’s sPG synthesis activity or FtsZ’s treadmilling speed can shift the partitioning of the synthesis complex between the two tracks, revealing a mechanism to coordinate sPG synthesis spatiotemporally with septal pole morphogenesis^15, 21, 23^. Such a two-track behavior was also observed in the diderm bacterium *C. crescentus*^24^. However, in Gram-positive (monoderm) bacteria such as *S. pneumoniae*^25^, *B. subtilis*^26^, and *S. aureus*^27^, only the slow-moving, sPG synthesis-driven population of synthases was observed. This difference likely suggests a less stringent regulation requirement for septation and cell pole morphogenesis due to the thicker cell walls in these bacteria, and/or a transitory Z-track-dependent behavior of these enzymes^28^.

Recently, we discovered that FtsN exhibits a single, slow-moving population on the sPG-track and is a required component of the active processive synthesis complex^21, 23^. Structural modeling shows that FtsN’s E domain binds to FtsL and FtsI in the previously identified Activation of FtsW and FtsI (AWI) domain^29^ and Constriction Control Domain (CCD)^29–31^, triggering a conformational change of FtsI’s Anchor domain to open up the catalytic pore of FtsW^15^. We also discovered that the moving dynamics of FtsN depend on its domain composition. For example, the isolated E domain in the absence of the rest of FtsN exhibits both Z-track and sPG-track populations and is sufficient to maintain the processivity of the synthesis complex. The cytoplasmic tail and the transmembrane domain of FtsN (FtsN^Cyto-TM^) exhibits only the Z-track dependent, fast-moving population^23^.

Besides the sPG-track-coupled slow-moving population, we observed that more than half of septal FtsN molecules are stationary because they are anchored to dnG by the SPOR domain^23^. While the essential E domain is the only region of FtsN strictly required for viability^9, 12^, the SPOR domain improves FtsN’s function by concentrating FtsN at the septum through its binding to dnG, which increases the rate of constriction and sPG synthesis^12, 32, 33^. FtsN also regulates sPG degradation by recruiting PG amidases that create dnG and mediate daughter cell separation^34^. Conversely, overproduction of FtsN’s SPOR domain impairs turnover of dnG by LTs^35^. Thus, the coexistence within one single protein of a SPOR domain that binds to dnG (a transient intermediate of sPG degradation) along with an E domain that activates the sPG synthesis complex, suggests that FtsN could link sPG synthesis and degradation in space and time^19, 36^.

In this work, we built on these ideas and dissected the coordination mechanism using SMT of FtsN and FtsW across various genetic backgrounds and growth conditions. Our studies identify a third track, termed as the dnG-track, characterized by the stationary population of FtsN bound to dnG via its SPOR domain. We discovered that FtsN’s SPOR domain self-interacts, binds to dnG cooperatively, and facilitates the formation of a multimeric sPG synthesis complex FtsWIQLB-FtsN on both the dnG- and sPG-tracks. Our results support a model in which FtsN, through the cooperative self-interaction of its SPOR domain, sensitively partitions between the dnG- and sPG-tracks. This partitioning regulates the balance between sequestered and active sPG synthesis complex, thereby coordinating sPG synthesis and degradation activities both in space and time.

## Results

### Altered dnG levels in Δ*amiABC* and Δ6*LT*s cells diminish sPG synthesis activity

To investigate the effect of SPOR-dnG binding on sPG synthesis activities, we monitored the moving dynamics of FtsN and FtsW using two mutant strains with altered dnG levels. In the Δ*amiABC* strain (strain EC5345, Supplementary Table 1), the dnG level is effectively zero due to the lack of all three periplasmic amidases^12, 19^, which generate dnG^7^. In the Δ6*LT*s strain (strain JL435, Supplementary Table 1), the dnG level is ∼20-40 fold higher than that in the wild type (WT) background^19^. Cells likely have long stretches of dnG strands due to the deletion of six out of eight known and putative LTs (Δ*mltA*, Δ*mltC*, Δ*mltD*, Δ*mltE*, Δ*slt70*, Δ*rlpA*), which degrade dnG^6^. Both the Δ*amiABC* and Δ6*LT*s mutants exhibited sick chaining phenotypes due to impaired ability to degrade sPG for proper daughter cell separation^6^ (Fig. 1a, b).

**Fig. 1.**
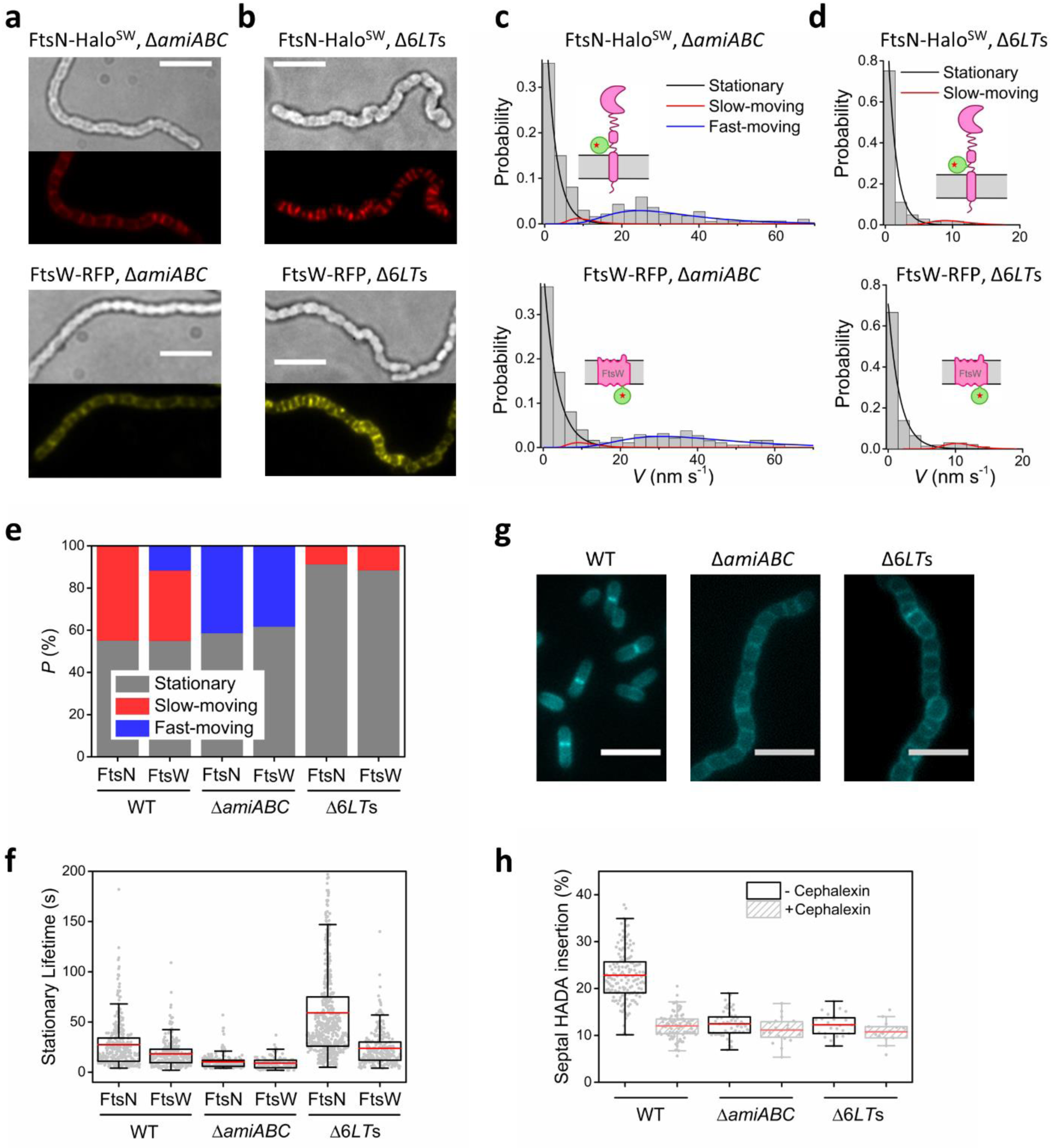
Altered dnG levels in Δ*amiABC* and Δ6*LT*s cells diminish sPG synthesis activity. **a-b**, Bright-field and ensemble fluorescent images of FtsN-Halo^SW^ (top) and FtsW-RFP (bottom) fusions in Δ*amiABC* (**a**) and Δ6*LT*s (**b**) strain backgrounds. Scale bars, 5 µm. **c-d**, Speed distributions of FtsN-Halo^SW^ (top) and FtsW-RFP (bottom) in Δ*amiABC* (**c**) and Δ6*LT*s (**d**) strain backgrounds overlaid with the fit curves of the stationary (black), slow-moving (red), and fast-moving (blue) populations. Schematic representation of the fusion and strain backgrounds are shown in inset (pink: labeled protein; green: Halo or RFP fluorophore). **e**, Percentages of the stationary (gray), slow-moving (red), and fast-moving (blue) populations of FtsN-Halo^SW^ and FtsW-RFP fusions in WT, Δ*amiABC*, and Δ6*LT*s strain backgrounds. **f**, Box plots of the lifetime of the stationary population under all the conditions in (**e**). Red lines indicate the mean values. **g**, Representative images of HADA-labeled WT, Δ*amiABC*, and Δ6*LT*s cells. Scale bars, 5 µm. **h**, Box plots of septal HADA fluorescence intensity percentage normalized to the whole cell intensity in WT, Δ*amiABC*, and Δ6*LT*s cells treated or untreated with cephalexin to inhibit FtsI.

To perform SMT of FtsN and FtsW, we ectopically expressed a functional, previously characterized FtsN-Halo^SW^ ^23^ or FtsW-TagRFP-t (abbreviated as FtsW-RFP)^21^ fusion protein from a plasmid in the two strain backgrounds (strains EC5345 and EC3708 respectively, Supplementary Table 1). As expected, and in accordance with altered dnG levels in the two strains, septal localization of FtsN-Halo^SW^ was reduced in Δ*amiABC* cells but enhanced in Δ6*LT*s cells (Fig. 1a, b, top; Supplementary Fig. 1a). FtsW-RFP showed similar localization levels (Fig. 1a, b, bottom; Supplementary Fig. 1b).

SMT of FtsN-Halo^SW^ and FtsW-RFP revealed that in Δ*amiABC* cells the two fusion proteins behaved similarly. The active, slow-moving populations on the sPG-track for both fusions were almost completely abolished (Fig. 1c, red curves). A new, fast-moving population of FtsN-Halo^SW^ appeared (*P*_fm_ = 41 ± 2%, *n* = 77 segments for FtsN-Halo^SW^), which was similar to that of FtsW-RFP (*P*_fm_ *=* 38 ± 3%, *n* = 72 segments, Fig. 1c-e, blue curves and bars, Supplementary Table 4). The average lifetime of stationary FtsN-Halo^SW^ molecules decreased to *T*_stat_ = 9.4 ± 0.6 s (*P*_stat_ = 59 ± 2%, *n* = 109 segments, Fig. 1f, Supplementary Table 4) from *T*_stat_ = 27 ± 1 s we previously measured under the WT condition^23^, comparable to that of stationary FtsW-RFP molecules (*T*_stat_ = 9.1 ± 0.6 s, *n* = 116 segments, Fig. 1f, Supplementary Table 4). This shortened stationary lifetime is similar to the average lifetime of stationary FtsZ molecules (8-12 s) measured in *E. coli* and other species^23, 25, 37–39^. As there is essentially no dnG in Δ*amiABC* cells, these results suggest that stationary FtsN-Halo^SW^ and FtsW-RFP molecules are stuck in the middle of treadmilling FtsZ polymers, as previously observed both *in vivo* and *in vitro*^21, 23, 40^.

In Δ6*LT*s cells, FtsN-Halo^SW^ and FtsW-RFP also behaved similarly to each other. The majority of the fusion protein molecules were stationary (*P*_stat_ = 91 ± 3%, *n* = 546 segments for FtsN-Halo^SW^ and *P*_stat_ = 88 ± 3%, *n* = 252 segments for FtsW-RFP, Fig. 1d, e, black curves and gray bars, Supplementary Table 4), consistent with the expectation that an elevated dnG level increases the number of dnG-bound FtsN-Halo^SW^ molecules and consequently FtsN-bound FtsW-RFP molecules. Most strikingly, the stationary lifetime of FtsN-Halo^SW^, which reflects the time it takes for single FtsN-Halo^SW^ molecules to dissociate from dnG, increased to *T*_stat_ = 59 ± 2 s in Δ6*LT*s cells (Fig. 1f, Supplementary Table 4). As we describe further below, the marked increase in both the percentage and dissociation time of stationary FtsN-Halo^SW^ molecules in Δ6*LT*s cells is most consistent with a higher SPOR-dnG affinity arising from cooperative binding of FtsN to long stretches of dnG. Similar phenomena have been observed on the length-dependent binding of oligomeric DNA- or mRNA-binding proteins^41–43^. The stationary lifetime of FtsW-RFP molecules also increased to *T*_stat_ = 24 ± 1 s (Fig. 1f, Supplementary Table 4), up from ∼18 s in the WT condition we measured previously^21^, suggesting FtsW may be bound cooperatively to multiple molecules of dnG-anchored FtsN (also see below).

The lack of slow-moving population of FtsN-Halo^SW^ and FtsW-RFP in the two strains indicates the lack of active sPG synthesis^21, 23^. To verify this result, we used HADA (HCC-Amino-D-alanine hydrochloride) labeling to measure the septal cell wall synthesis activity^22^. Both mutants showed significantly lower HADA incorporation levels than WT cells, which were indistinguishable from sPG synthesis-inhibited cells treated with cephalexin (Fig. 1g, h). The lack of septal cell wall synthesis activity in Δ*amiABC* cells is consistent with the minimal septal localization of FtsN in the absence of dnG (Fig.1a). However, in Δ6*LT*s cells where septal FtsN and FtsW are abundant, the lack of HADA incorporation implies that the FtsWIQLB-FtsN complex is sequestered on dnG and needs to be released from dnG to engage in sPG synthesis.

### Truncation of FtsN’s SPOR domain leads to the sequestering of FtsW on the Z-track

To examine the role of FtsN-dnG interaction without modulating dnG levels, we constructed an *ftsN*^ΔSPOR^ strain in which the *spor* sequence is deleted from the full-length *ftsN* gene at its native chromosomal locus (strain EC3338, Supplementary Table 1). Cells expressing the *ftsN*^ΔSPOR^ truncation gene grew slowly and were filamentous, but expression of an additional copy of *ftsN*^ΔSPOR^ from a plasmid restored WT-like growth and similar septum synthesis activity (strain JL337, termed as *ftsN*^ΔSPOR+^ herein for simplicity, Supplementary Table 1, Supplementary Fig. 2). These results are consistent with the fact that SPOR is the major septum localization determinant of FtsN, and hence a higher expression level of FtsN^ΔSPOR^ is needed to support cell division at a similar level^12^.

Next, we performed SMT of FtsW-RFP expressed from a plasmid in the *ftsN*^ΔSPOR+^ background (strain JL362, Supplementary Table 1) using the same M9 minimal medium and imaging conditions as before^23^. We observed that the sPG synthesis processivity of FtsW, defined as the persistent running time on the sPG-track, remained similar in the absence of SPOR (*T*_sm_ = 21.1 ± 0.5 s, *μ* ± s.e.m. in *ftsN*^ΔSPOR+^ *vs.* 20.1 ± 0.1 s in WT, Fig. 2b, top, Supplementary Fig. 3, Supplementary Table 5). The Z-track processivity of FtsW, similarly defined as the persistent running time on the Z-track, remained unchanged as well (*T*_fm_ = 9.4 ± 0.2 s, *μ* ± s.e.m. in *ftsN*^ΔSPOR+^ *v.s.* 8.3 ± 0.3 s in WT, Fig. 2b, bottom, Supplementary Table 5). However, the active, slow-moving population of FtsW-RFP decreased significantly (from 33 ± 2%, *n* = 157 segments in WT to 8 ± 3%, *n* = 38 segments in *ftsN*^ΔSPOR+^), while the fast-moving population increased from 12 ± 2% to 19 ± 2% (Fig. 2a, c, Supplementary Table 5). Additionally, the stationary population of FtsW-RFP increased to 73.5 ± 2% (*n* = 353 segments, Fig. 2c, Supplementary Table 5) from ∼55% under the WT condition as we previously reported^21^ and had a short lifetime of *T*_stat_ = 10.4 ± 0.4 s (Supplementary Table 5). This stationary lifetime is comparable to that of FtsW-RFP in Δ*amiABC* cells (Supplementary Table 4), suggesting that they were similarly stuck in the middle of FtsZ polymers. Taken together, these results suggest that in the absence of FtsN’s SPOR domain, while FtsW’s processivities on the sPG- and Z-tracks are largely unaltered, FtsW has a significantly reduced active, slow-moving population on the sPG-track and is primarily sequestered on the Z-track.

**Fig. 2.**
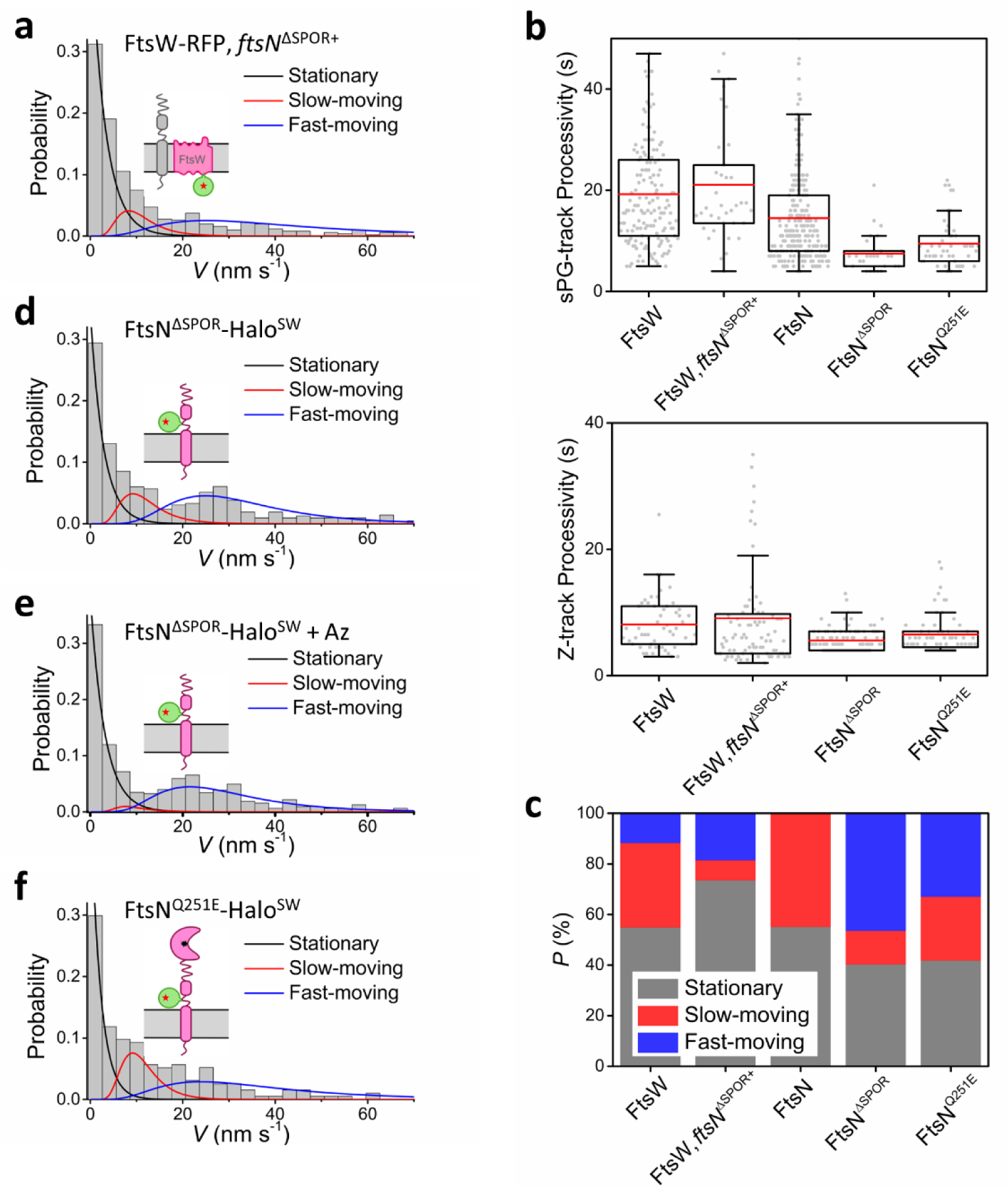
SPOR domain enhances FtsW and FtsN’s slow-moving populations. **a**, **d-f**: Speed distributions of FtsW-RFP in *ftsN*^ΔSPOR+^ strain background (**a**), FtsN^ΔSPOR^-Halo^SW^ in *ftsN*^ΔSPOR+^ strain background (**d**) and treated with aztreonam (**e**), and FtsN^Q251E^-Halo^SW^ in WT background (**f**), overlaid with the fit curves of the stationary (black), slow-moving (red), and fast-moving (blue) populations. Schematic representation of the fusion and strain background are shown in insets. **b**, sPG-track processivity (top; defined as the persistent running time of the slow-moving population) and Z-track processivity (bottom; defined as the persistent running time of the fast-moving population) under the conditions in (**a**, **d-f**). **c**, Percentages of the stationary (gray), slow-moving (red), and fast-moving (blue) populations under all the conditions in (**a**, **d-f**). FtsW and FtsN data were replotted from figure (**1e**) for comparison.

### Truncation of FtsN’s SPOR domain leads to the sequestering of FtsN^ΔSPOR^ on the Z-track and reduces its sPG-track processivity

To examine how truncation of the SPOR domain affects the moving dynamics of FtsN itself, we constructed a chromosomal integrated FtsN^ΔSPOR^-Halo^SW^ fusion protein and overexpressed it in an *ftsN-*depletion background (strain EC5257, Supplementary Table 1) to restore normal cell length. We observed that the stationary population of FtsN^ΔSPOR^-Halo^SW^ at septa decreased to 40% ± 2% with an average *T*_stat_ = 9.2 ± 0.6 s (*n* = 85 segments, Fig. 2c, d, Supplementary Table 5), suggesting that they were trapped in the middle of FtsZ polymers. Most importantly, in contrast to the single slow-moving population observed for FtsN-Halo^SW^, FtsN^ΔSPOR^-Halo^SW^ exhibited two distinct processively-moving populations, one slow (*P*_sm_ = 13 ± 3%, *V*_sm_ = 9.4 ± 0.1 nm s^-1^, *μ* ± s.e.m., *n* = 28 segments) and one fast (*P*_fm_ = 46 ± 2%, *V*_fm_ = 32.7 ± 0.1 nm s^-1^, *μ* ± s.e.m., *n* = 98 segments, Fig. 2c, d, Supplementary Table 5). Inhibiting sPG synthesis activity using aztreonam completely abolished the slow-moving population of FtsN^ΔSPOR^-Halo^SW^ but preserved the fast-moving population (Fig. 2e, Supplementary Table 5), similar to what we previously observed for FtsW in aztreonam-treated cells^21^ or FtsB in fosfomycin-treated cells^15^. However, the persistent running times, or the processivities, of FtsN^ΔSPOR^-Halo^SW^ on both the sPG-track and the Z-track (*T*_sm_ = 7.5 ± 0.1 s and *T*_fm_ = 5.5 ± 0.1 s respectively, Fig. 2b, Supplementary Table 5) were shorter than those of FtsW-RFP in the *ftsN*^ΔSPOR+^ background and full-length FtsN-Halo^SW^ on the sPG-track (14.5 ± 0.7 s, Fig. 2b, Supplementary Table 5). These results suggest that similar to FtsW in the absence of FtsN’s SPOR domain, septal FtsN^ΔSPOR^-Halo^SW^ was mainly sequestered on the Z-track. Because dnG-bound FtsN molecules are stationary and not running on the sPG-track^23^, the reduced persistent running time of FtsN^ΔSPOR^-Halo^SW^ on the sPG-track suggests that the SPOR domain may have a role in supporting the processivity of FtsN in the active sPG synthesis complex, independent of its dnG-binding ability.

### An FtsN^Q251E^ mutant defective in dnG-binding partially restores FtsN’s sPG-track population and processivity

Truncation of FtsN’s SPOR domain removes both its dnG-binding ability and potentially other interactions of the SPOR domain independent of dnG binding. To further investigate the effects of these two types of interactions, we monitored the moving dynamics of an FtsN^Q251E^-Halo^SW^ fusion protein by transforming a plasmid expressing *ftsN*^Q251E^*-halo*^SW^ into an *ftsN*-depletion strain (strain JL378, Supplementary Table 1). Residue Q251 is the most highly conserved amino acid in the dnG-binding cleft and interacts directly with the glycan chain^16^. The Q251E substitution reduces septal localization and dnG-binding of the isolated SPOR domain but has not been studied in the context of full-length FtsN^44^.

Consistent with a defect in dnG-binding, FtsN^Q251E^-Halo^SW^ exhibited a fast-moving population on the Z-track (*P*_fm_ = 33 ± 2%, *T*_fm_ = 6.4 ± 0.2 s, *n* = 64 segments, Fig. 2b, f, Supplementary Table 5) and a smaller, 42 ± 2% stationary population with a significantly reduced stationary lifetime of *T*_stat_ = 13.1 ± 0.9 s (*n* = 81 segments, Fig. 2c, Supplementary Table 5). These results further confirm that the dnG-binding ability of SPOR prevents the release of FtsN to the Z-track. Interestingly, the slow-moving population percentage and processivity of FtsN^Q251E^-Halo^SW^ recovered to *P*_sm_ = 25 ± 2% and *T*_sm_ = 9.7 ± 0.2 s (*n* = 49 segments), in between that of FtsN^ΔSPOR^-Halo^SW^ (*P*_sm_ = 13% and *T*_sm_ = 7.5 s) and full-length FtsN-Halo^SW^ (*P*_sm_ = 45% and *T*_sm_ = 14.5 s) (Fig. 2b, c, Supplementary Tables 4 and 5). These results suggest that both the dnG-binding ability and the dnG-binding-independent interaction of SPOR contribute to maintaining the active sPG synthesis complex on the sPG-track (also see **Discussion**).

### Isolated SPOR domain (iSPOR) moves processively with WT FtsN

To further investigate the role of the SPOR domain in supporting FtsN’s slow-moving population on the sPG-track independent of FtsN’s other domains, we used a previously characterized fusion construct FtsN^WYAA^-Halo^SW^ ^23^. This fusion protein contains two critical mutations, W83A and Y85A, in FtsN’s E domain, which abolish the binding of the E domain to FtsI and FtsL and the subsequent activation of FtsWI^9, 45^. We previously reported that FtsN^WYAA^-Halo^SW^ molecules were completely stationary at septa in the superfission variant *ftsB*^E56A^ Δ*ftsN* background^23^, despite that FtsW-RFP still moved processively on the sPG-track in the same strain background^21^.

Surprisingly, when we ectopically expressed this FtsN^WYAA^-Halo^SW^ fusion protein in the presence of WT FtsN (Fig. 3a, strain EC5259, Supplementary Table 1), it exhibited essentially the same percentage of the slow-moving population compared to WT FtsN-Halo^SW^ (*P*_sm_ = 44 ± 4%, *n* = 79 segments, Fig. 3b, c, Supplementary Table 6) with a modestly reduced processivity (*T*_sm_ = 10.7 ± 0.8 s, Fig. 3d, top, Supplementary Table 6). This result suggests that FtsN^WYAA^-Halo^SW^ remains part of the processively-moving sPG synthesis complex through the binding of domains other than the E domain to WT FtsN in the complex, as the direct binding of its E domain with FtsI and FtsL is abolished by the WYAA mutations.

**Fig. 3.**
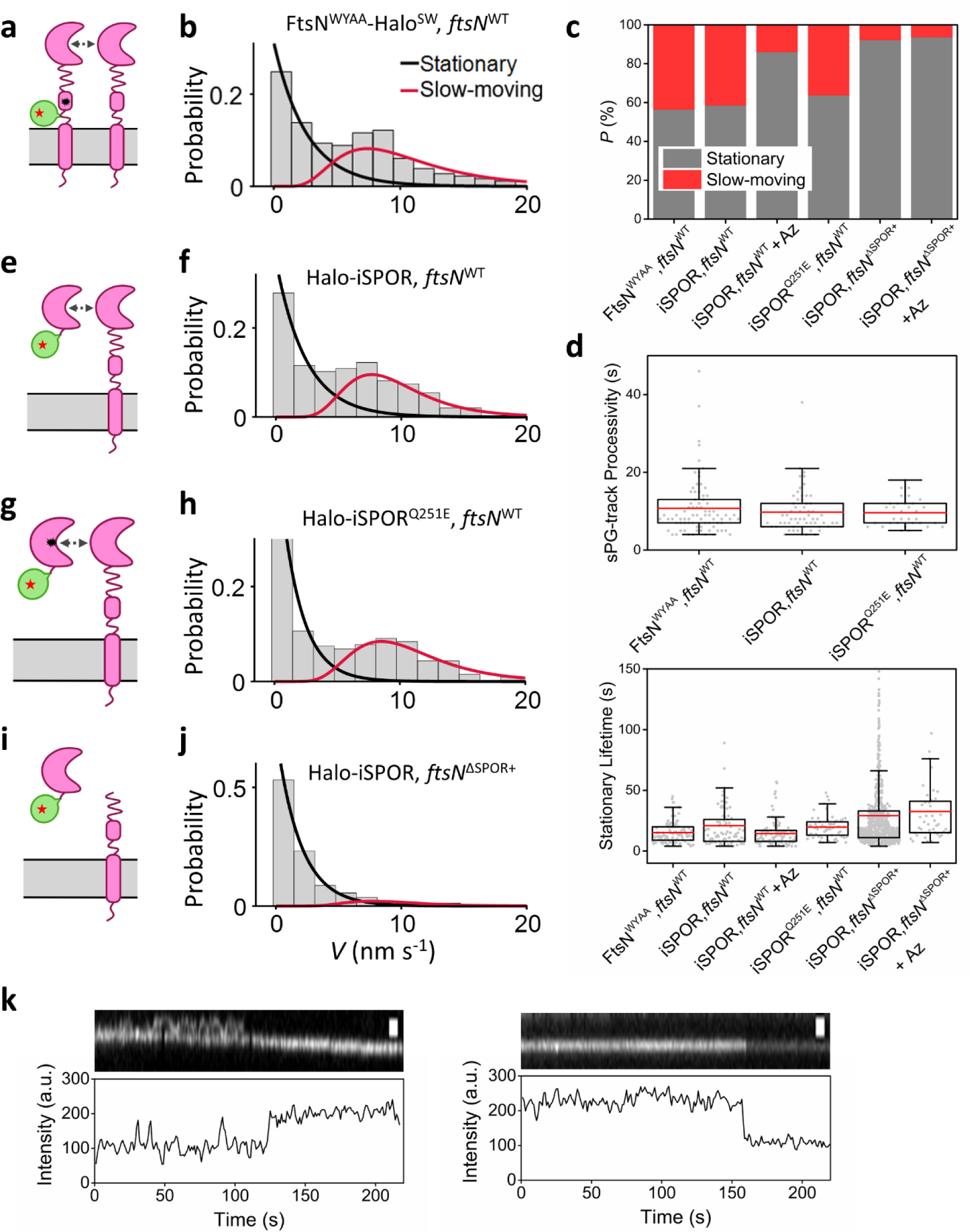
SPOR domain exhibits SPOR-SPOR self-interaction. **a**, Schematic representation of FtsN^WYAA^-Halo^SW^ fusion in WT *ftsN* background. **b**, speed distribution of FtsN^WYAA^-Halo^SW^ overlaid with the fit curves of the stationary (black) and slow-moving (red) populations. **c**, Percentages of the stationary (gray) and slow-moving (red) populations under all the conditions in **b**, **f**, **h**, **j**, and **S3**. **d**, Top: sPG-track processivity of FtsN^WYAA^-Halo^SW^, Halo-iSPOR, and Halo-iSPOR^Q251E^ fusions in WT strain. Bottom: Stationary lifetime of all the conditions in (**c**). **e**, Schematic representation of Halo-iSPOR fusion in WT *ftsN* background. **f**, speed distribution of Halo-iSPOR overlaid with corresponding fit curves. **g**, Schematic representation of Halo-iSPOR^Q251E^ fusion in WT *ftsN* background. **h**, Speed distribution of Halo-iSPOR^Q251E^ overlaid with corresponding fit curves. **i**, Schematic representation of Halo-iSPOR fusion in *ftsN*^ΔSPOR+^ strain background; **j**, speed distribution of Halo-iSPOR overlaid with corresponding fit curves. **k**, Representative kymographs (top) and intensity time traces (bottom) of FtsN-Halo^SW^ multimers at the septum. Left: an FtsN-Halo^SW^ molecule transiently associated and disassociated (diffusive fluorescence intensity) with a processively moving FtsN-Halo^SW^ molecule (stable fluorescence intensity) between 20 s and 100 s, and eventually stably associated (sudden doubling of fluorescence intensity) and moved together at an average speed of 7 nm s^-1^; Right: one FtsN-Halo^SW^ molecule was photo-bleached (∼155 s) in a stationary dimer. Scale bars, 1 µm.

Our previous studies showed that the cytoplasmic and transmembrane domains of FtsN are not involved in the formation of the processive sPG complex^23^. Therefore, we reasoned that the SPOR domain might be responsible for such interaction in the absence of the E domain-mediated binding to FtsI and FtsL. We constructed a Halo-iSPOR fusion protein, in which a Halo-tag is fused to the N-terminus of the isolated SPOR domain (hereafter termed as iSPOR to distinguish from the SPOR domain linked to the rest of FtsN), and expressed it in the presence of WT FtsN (Fig. 3e, strain EC5275, Supplementary Table 1). Strikingly, while Halo-iSPOR localized to the septum as expected (Supplementary Fig. 4), these Halo-iSPOR molecules moved directionally (*P*_sm_ = 42 ± 3%, *v* = 9.4 ± 0.5 nm/s, *n* = 61 segments, Fig. 3c, f, Supplementary Table 6) with a processivity of *T*_sm_ = 9.8 ± 0.7 s (Fig. 3d, top, Supplementary Table 6), and became predominately stationary in cells treated with aztreonam (Supplementary Fig. 5), mimicking what we observed for full-length FtsN-Halo^SW^ previously^23^. These results strongly suggest that there exists SPOR-SPOR self-interaction.

### iSPOR^Q251E^ defective in dnG-binding moves processively with WT FtsN

An alternative explanation for the observed processive movement of iSPOR with WT FtsN is that iSPOR and FtsN’s SPOR domain may bind to the same dnG fragment disintegrated from the cell wall without direct protein-protein interactions. To examine this possibility, we constructed an isolated Halo-iSPOR^Q251E^ fusion, which is defective in dnG-binding, and expressed it in the WT *ftsN* background (Fig. 3g, strain JL431, Supplementary Table 1). Septal localization of Halo-iSPOR^Q251E^ decreased about 2-fold compared to unmutated Halo-iSPOR (Supplementary Fig. 4b). Surprisingly, Halo-iSPOR^Q251E^ still exhibited a large slow-moving population with a similar processivity compared to Halo-iSPOR (*P*_sm_ = 36 ± 4%, *T*_sm_ = 9.6 ± 0.6 s, *n* = 33 segments, Fig. 3c, d, h, Supplementary Table 6). The apparent ∼2-fold decrease in dnG-binding with little or no change in the dynamics of the slow-moving population argues the observed iSPOR-FtsN interaction is independent of dnG-binding. Note that the moving dynamics of Halo-iSPOR^Q251E^ in the presence of WT FtsN mimics that of WT FtsN but not that of FtsN^Q251E^-Halo^SW^. The latter exhibited a fast-moving population (compare Fig. 2f and Fig. 3h) because it has an N-terminal cytoplasmic domain that can bind to FtsA, whereas Halo-iSPOR^Q251E^ does not. This comparison further confirms that there must exist SPOR-SPOR self-interaction in the processive sPG synthesis complex.

### iSPOR becomes stationary when disconnected from the rest of FtsN

To further verify that iSPOR binds to the existing SPOR domain of full-length FtsN in the processive sPG synthesis complex, we expressed Halo-iSPOR *in trans* with FtsN^ΔSPOR+^ (Fig. 3i, strain EC5415, which also contains an extra plasmid copy of *ftsN*^ΔSPOR^, Supplementary Table 1). Halo-iSPOR localized to the septum in the *ftsN*^ΔSPOR+^ background similarly as in the WT *ftsN* background (Supplementary Fig. 4). We reasoned that if the movement of iSPOR were due to its binding to FtsN domains other than SPOR, we should still observe its processive movement with FtsN^ΔSPOR+^. However, Halo-iSPOR molecules became completely stationary (Fig. 3c, j), with a stationary lifetime of *T*_stat_ = 29 ± 1 s (*n* = 602 segments, Supplementary Table 6) that is indistinguishable from full-length FtsN-Halo^SW^ (Fig. 1f). Treating cells with aztreonam did not change the percentage nor the lifetime of these stationary Halo-iSPOR molecules (Fig. 3c, d, Supplementary Fig. 5). Because we showed that in the same *ftsN*^ΔSPOR*+*^ background, FtsW-RFP and FtsN^ΔSPOR^-Halo^SW^ exhibited both slow- and fast-moving populations (Fig. 2), these results strongly suggest that SPOR-SPOR self-interaction and the linkage between SPOR and the rest of FtsN are responsible for the observed processive movement of iSPOR. As such, these results demonstrate that the active, processive FtsWIQLB-FtsN complex contains multiple FtsN molecules mediated by SPOR-SPOR self-interaction, and that this interaction is important to sustain the active population of sPG synthesis complex on the sPG-track.

We also note that the stationary lifetimes (∼15-20 s) of the FtsN^WYAA^-Halo^SW^, Halo-iSPOR and Halo-iSPOR^Q251E^ in the presence of WT FtsN were shorter than that of full-length FtsN-Halo^SW^ (∼30 s, Fig. 3d, bottom, Supplementary Table 6), suggesting that FtsN^WYAA^-Halo^SW^ and iSPOR domains may not compete effectively with WT FtsN for dnG-binding. Interestingly, in ensemble time-lapse movies of cells treated with aztreonam, FtsN-Halo^SW^ signal also persisted at the septum longer than either Halo-iSPOR or FtsN^ΔSPOR^-Halo^SW^ (Supplementary Fig. 6, Supplementary Movies 1-3). The longer persistence of full-length FtsN-Halo^SW^ compared to Halo-iSPOR or FtsN^ΔSPOR^-Halo^SW^ further suggest that the SPOR domain must be linked to the rest of FtsN in *<u>cis</u>* in the same molecule for the full functionality of FtsN.

### Stationary and processive sPG synthesis complexes are non-monomeric

In our single-molecule tracking experiments, we used sparse Halo-dye (JF646) labeling to ensure the tracking of single molecules of interest. We also screened out fluorescent spots with intensities higher than that of single molecules. In light of the above results, we reexamined some of our data to assess whether we could directly visualize the presence of more than one molecule in the same stationary and processive sPG synthesis complex. Indeed, we identified FtsN-Halo^SW^ molecules in both moving and stationary SMT trajectories where the fluorescent spot intensities are significantly higher than expected from a single JF646 dye molecule, which also exhibited the typical quantized photobleaching step or fluorescence jump for single molecules (Fig. 3k; Supplementary Movies 4-5). We even observed one FtsN-Halo^SW^ molecule that repeatedly associated and dissociated until it stably associated with a processively moving complex (Fig. 3k, left; Supplementary Movie 4). Most importantly, we identified similar trajectories showing the presence of more than one single molecule from our previous SMT of Halo-FtsB^15^, FtsI-Halo^SW^ ^20^ and FtsW-RFP^21^ fusions in *E. coli* and Halo-FtsW in *C. crescentus*^24^ (Supplementary Fig. 7; Supplementary Movies 6-8). Thus, while the sparse labeling prevents us from further statistically analyzing the number of molecules in the complex, these observations demonstrate that the FtsWIQLB-FtsN complex in *E. coli* exists as a multimer in the form of (FtsWIQLB)_m_-FtsN_n_ or (FtsWIQLB-FtsN)_n_. The non-monomeric nature of the sPG synthesis complex may be a conserved feature across different bacterial species, as a recent SMT study in *B. subtilis* also reported the presence of multimeric PBP2b, the equivalent of FtsI in *E. coli*, on the sPG-track^26^.

### Relative sPG synthesis and degradation activities are correlated in WT *ftsN* background but skewed when SPOR is disconnected

Under normal growth conditions in *E. coli*, sPG synthesis and degradation activities are balanced to avoid aberrant cell pole morphogenesis or septal cell wall lesions. FtsN has been proposed as a key player in coordinating these two opposing activities by linking sPG synthesis to the abundance and location of dnGs^19, 36^. It is known that the binding of FtsN’s E domain to the FtsWIQLB complex activates sPG synthesis activity^15, 23, 46–48^. We also further confirmed that overexpressing iSPOR in WT cells caused a profound chaining phenotype (Supplementary Fig. 8), suggesting that the binding of FtsN’s SPOR domain to dnG protects them from being degraded by LTs, as also suggested by a previous study^35^. Therefore, we propose that the partitioning of FtsN between the two populations, one slow-moving in complex with active FtsWIQLB and the other stationary bound to dnG, would naturally balance sPG synthesis and degradation activities: release of FtsN and FtsN-bound FtsWIQLB complex from dnG leads to processive sPG synthesis while simultaneously exposing dnGs to degradation by LTs, enhancing both synthesis and degradation; conversely, binding of FtsN to dnG protects dnG and also sequesters FtsN-bound FtsWIQLB complex, diminishing both synthesis and degradation.

To analyze the proposed correlation quantitatively, we compared FtsN’s occupancies on the processive sPG-track and dnGs (Fig. 4). We calculated FtsN’s occupancy on the sPG-track (*N*_sPG_) using the product of the slow-moving population percentage of FtsN-Halo^SW^ and the corresponding persistent running time and velocity under each condition, normalized to that of the WT condition (**Methods**). Note that FtsN’s occupancy on the sPG-track is highly correlated with that calculated using FtsW-RFP data (Supplementary Fig. 9). Additionally, we previously showed that FtsW’s occupancy on the processive sPG-track correlated highly to the cell wall constriction rate and HADA incorporation level^21^. Therefore, *N*_sPG_ can be used as a proxy for sPG synthesis activity.

**Fig. 4.**
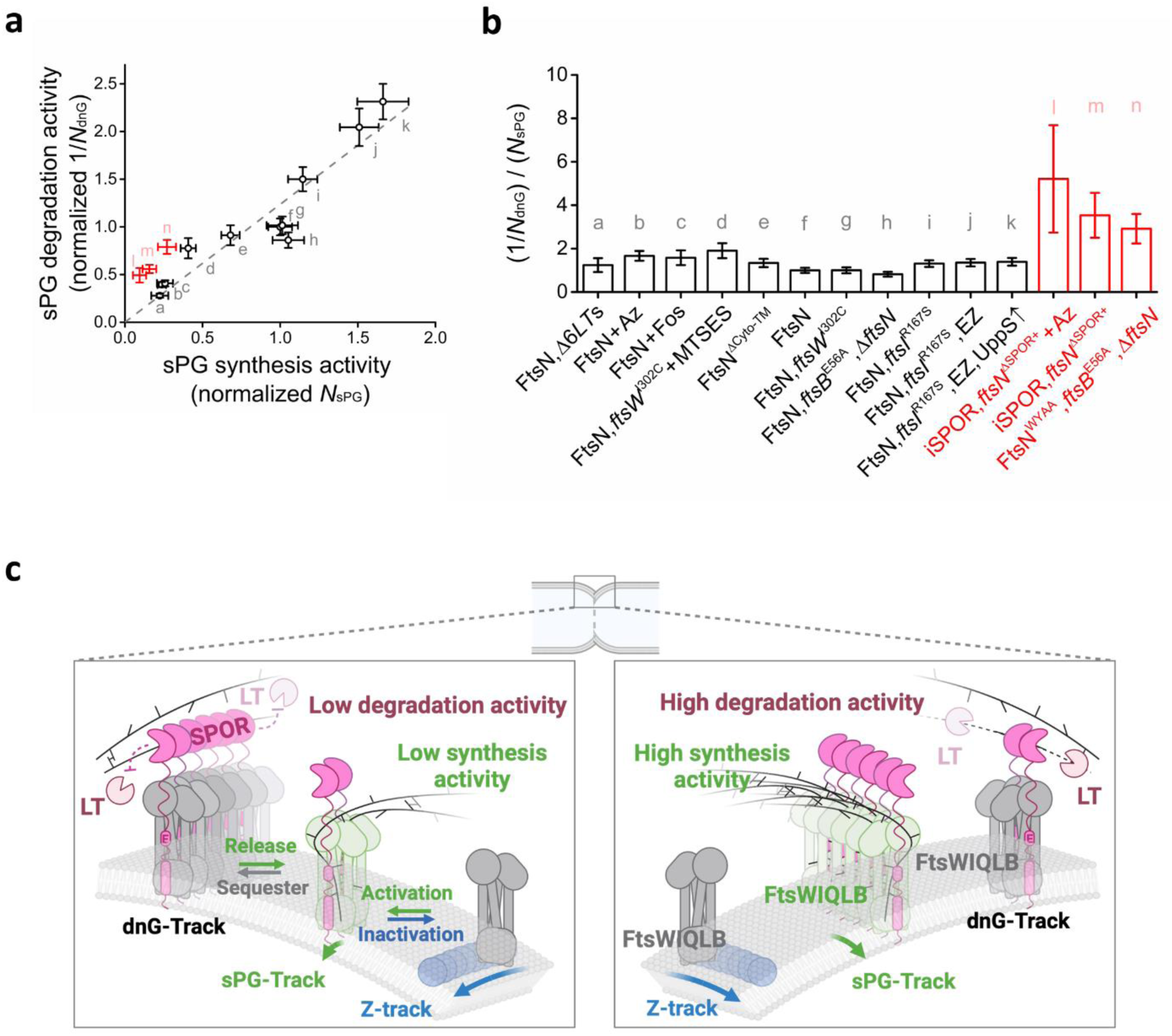
FtsN coordinates sPG synthesis and degradation both in space and time. **a**, The reverse of normalized FtsN occupancy on dnG (1/*N*_dnG_) as a proxy for sPG degradation activity correlates with normalized FtsN occupancy on sPG (*N*_sPG_) across all conditions (black), except when the SPOR domain is unlinked from the rest of the FtsN (red). All the data points are labeled with alphabetical letters, in the order of elevated sPG occupancy (*N*_sPG_) as a proxy for sPG synthesis activity. **b**, The ratio between 1/*N*_dnG_ and *N*_sPG_, the proxy for the ratio between relative sPG degradation and synthesis activities is close to one (black), but became significantly higher when the SPOR domain was unlinked from the rest of the FtsN (red). The conditions are labeled with the same letters in (**a**). **c**, A three-track model depicting how FtsN coordinates sPG synthesis and degradation both in space and time. FtsN marks a third track, termed as the dnG-track, through its SPOR domain’s binding to dnG. When sPG synthesis activity is low (e.g., under antibiotic treatment, left), FtsN shifts from the sPG-track to the dnG-track and consequently sequesters FtsWIQLB on dnG. Cooperative binding of SPOR to dnG inhibits LTs’ access to dnG, thus decreasing sPG degradation activity correspondingly. Conversely, when sPG synthesis activity is high (e.g., in rich medium, right), FtsN shifts from the dnG-track to the sPG-track, thereby exposing dnG and facilitating its degradation by LTs, consequently increasing sPG degradation activity.

We calculated FtsN’s occupancy on dnG (*N*_dnG_) using the product of the percentage of the stationary population of FtsN-Halo^SW^ and its corresponding stationary lifetime, normalized to that of the WT condition (**Methods**). As *N*_dnG_ reflects the level of dnG protected by FtsN, the inverse of *N*_dnG_ (1/*N*_dnG_) should reflect LTs’ sPG degradation activity (high *N*_dnG_ indicates low LT activities and low *N*_dnG_ indicates high LT activities).

In Fig. 4a, we plotted 1/*N*_dnG_ (proxy for LTs’ sPG degradation activity) against *N*_sPG_ (proxy for sPG synthesis activity) and observed that 1/*N*_dnG_ is highly correlated with *N*_sPG_ across a variety of conditions (*p*_pearson_ = 0.9412). For example, in cells harboring a superfission variant FtsI^R167S^ and grown in rich medium, both *N*_sPG_ and 1/*N*_dnG_ were high, indicating high activities in both sPG synthesis and degradation (Fig. 4a, conditions j, k). In cells treated with cell wall synthesis inhibitors (MTSES for *ftsW*^I302C^, antibiotics aztreonam or fosfomycin) or deleted of six LTs (Δ6*LT*s), both *N*_sPG_ and 1/*N*_dnG_ were low, indicating low activities in both sPG synthesis and degradation (Fig. 4a, conditions a-d). A modified FtsN variant that lacks the cytoplasmic and transmembrane domains (Halo-FtsN^ΔCyto-TM^) also largely followed the same trend (Fig. 4a, condition e), indicating that the correlation is unrelated to the cytoplasmic interactions of FtsN with FtsA.

Most strikingly, we found that on average, the ratio between 1/*N*_dnG_ and *N*_sPG_, the proxy for the ratio between relative sPG degradation and synthesis activities, was close to one across all conditions that contained a functional, full periplasmic copy of FtsN (Fig. 4b, black bars), irrespective of whether sPG synthesis is enhanced (as in superfission backgrounds, conditions i-k) or inhibited (as in cells treated with antibiotics, conditions a-d). However, this ratio became significantly higher (Fig. 4a, b, red) at ∼3 when the E domain’s binding to FtsWIQLB was disabled by the WYAA mutations (condition n), or in cells expressing Halo-iSPOR in *trans* with *ftsN*^ΔSPOR+^ (condition m), and further increased to ∼5 when cells were treated with aztreonam (condition l). These results hence suggest that FtsN’s partitioning on the sPG- and dnG-tracks maintains balanced sPG synthesis and degradation activities, and that the linking of the SPOR domain to the rest of FtsN is required for this role.

## Discussion

In *E. coli*, sPG synthesis and turnover must be meticulously coordinated to prevent lysis and ensure synchronous invagination of the three layers of the envelope. The essential bitopic membrane protein FtsN was suggested to coordinate sPG synthesis and turnover because its periplasmic E domain activates the sPG synthesis complex FtsWIQLB and its C-terminal SPOR domain binds to dnG, a transient intermediate in biogenesis of the division septum. Here we investigated this role using SMT to monitor the dynamics of septal FtsN and FtsW under various conditions. Our findings yield several significant insights and lead to a model explaining how FtsN coordinates sPG synthesis and degradation in space and time, which we discuss below.

### A dnG-track anchors stationary FtsWIQLB-FtsN complexes

We previously reported the existence of stationary FtsW-RFP and FtsI-Halo^SW^ molecules with an average stationary lifetime of ∼18-20 s, which reflects the contributions of two populations of FtsWI: one with a short stationary lifetime (∼9 s) bound to internal positions in FtsZ polymers, and another with a longer stationary lifetime whose location was yet unknown^21, 23^. These longer-lived stationary synthase molecules were inferred as “poised” for sPG synthesis, as their abundance decreased under conditions stimulating sPG synthesis, such as overproduction of UppS to provide more Lipid II, and increased under conditions inhibiting sPG synthesis, such as treating cells with aztreonam or fosfomycin^21, 23^. Our current findings suggest that these “poised” FtsWI molecules are bound to dnG-anchored FtsN. This suggestion is supported by the disappearance of the long-lived FtsW-RFP stationary population in Δ*amiABC* cells, which lack dnG (average stationary lifetime reduced to 9 s, Fig. 1f, Table S4). Conversely, the average stationary lifetime of FtsW-RFP increased to ∼24 s in Δ6*LT*s mutant (Fig. 1f, Table S4), where dnG-anchored FtsN accumulates to high levels.

We refer to poised sPG synthases bound to dnG-anchored FtsN as being on the “dnG-track”. Along with the previously identified Z-track and sPG-track^15, 21, 23^, the dnG-track brings the total number of notional tracks in the *E. coli* septum to three. What are their purposes? We previously showed that the dynamic redistribution of FtsWIQLB between the Z-track and sPG-track facilitates the even distribution of enzymes along the septum, ensuring smooth and symmetric septum growth^21^. In contrast, the dnG-track likely functions to synchronize sPG synthesis and degradation both in space and time (see below).

### FtsN ensures balanced sPG synthesis and degradation activities

Our results show that under most conditions where FtsN is intact, the normalized activities of sPG synthesis and degradation are proportional, i.e., balanced. However, this balance is disrupted in cells where the SPOR domain (iSPOR) is disconnected from the rest of the FtsN (Fig. 4a, b, red). These findings align with a previous study using fluorescent D-amino acids (FDAAs) to compare septal cell wall synthesis and degradation rates in WT and an *ftsN*^ΔSPOR^ mutant^32^. That study found a clear decrease in sPG synthesis and a marginal increase in sPG degradation in *ftsN*^ΔSPOR^ cells, which was not considered significant but nevertheless trended in the expected direction of increased degradation/synthesis ratio if FtsN’s SPOR domain protects dnG from LT activity. The modest effect of *ftsN*^ΔSPOR^ on sPG degradation suggests other SPOR proteins may help protect dnG in the absence of FtsN’s SPOR domain. Interestingly, using FDAAs to assay sPG synthesis and turnover revealed a lower degradation/synthesis ratio in a superfission *ftsL*\* (*ftsL*^E88K^) background. This strain still contained a functional full-length FtsN. It would be interesting to examine whether FtsN’s binding to this mutant FtsL is increased, as the mutation FtsL^E88K^ is next to FtsL^E87^ that is involved in the binding with FtsN’s E domain^15^.

### FtsN coordinates sPG synthesis and degradation both in space and time

Our findings support a model in which FtsN not only coordinates sPG synthesis and degradation at the activity level (Fig. 4a, b), but also in space and time (Fig. 4c). Specifically, FtsN binds to dnG generated by amidases, protecting them from degradation by LTs. The sPG synthesis complex FtsWIQLB becomes anchored to dnG-bound FtsN via its E domain binding to FtsI and FtsL. When FtsN’s SPOR domain dissociates from dnG and when sPG precursors are available, the FtsWIQLB-FtsN complex initiates processive sPG synthesis in the vicinity of dnG. Simultaneously, the dissociation of FtsN’s SPOR domain from dnG allows LTs to destroy recently-vacated binding sites for FtsN. Similarly, when the FtsWIQLB-FtsN complex moves processively to synthesize new sPG, the long, flexible linker of FtsN connecting the SPOR and E domains may allow SPOR to probe for newly generated dnGs nearby. Once found, SPOR binds to these new dnGs, potentially bringing over the entire FtsWIQLB-FtsN complex, which will then commence another round of sPG synthesis nearby when released again. This iterative cycling of sPG synthesis and degradation ensures that these two activities are closely coordinated both in space and time.

### Self-interaction of SPOR

Our study reveals that FtsN’s SPOR domain self-interacts both when bound to dnG and when bound to the active FtsWIQLB complex. The following lines of evidence support this observation. First, FtsN is expressed at a level about 3-4-fold higher than FtsW, FtsI, and FtsQLB^23, 49^, making it possible to form at least (FtsWIQLB-FtsN)_n_ type of complexes. Second, our previous Structured Illumination Microscopy (SIM) imaging experiments showed large FtsN clusters that either remained stationary or moved processively together^23^. Third, in our current SMT experiments, we directly observed more than one FtsN molecule in the stationary or slow-moving trajectories within our typical imaging resolution limit of ∼30-40 nm^23^ (Fig. 3k). Fourth, FtsN^WYAA^ moved processively in the presence of full-length WT FtsN (Fig. 3a), but became completely stationary when *ftsN* was deleted in the superfission *ftsB^E56A^* background^23^. Similarly, iSPOR moved processively in the presence of full-length WT FtsN (Fig. 3e, f), but not in the presence of FtsN^ΔSPOR+^ (Fig. 3i, j). Finally, iSPOR^Q251E^, which is defective in dnG binding, still exhibited similar percentages of both slow-moving and stationary populations (Fig. 3g, h, Supplementary Table 4). Interestingly, the septal localization percentage of iSPOR^Q251E^ was half that of Halo-iSPOR, yet the percentage was similar in the presence and absence of full-length FtsN (Supplementary Fig. 4). This independence suggests that other SPOR proteins (of which *E. coli* has three^12, 50^) may substitute for FtsN’s SPOR domain. Taken together, these observations strongly support the presence of SPOR-SPOR self-interaction independent of dnG-binding.

We have tried to characterize the self-interaction of iSPOR *in vitro* but were largely unsuccessful. Purified iSPOR showed a single monomeric species in size exclusion columns and static light scattering (Supplementary Fig. 11, Supplementary Table 7). Crosslinking experiments to trap transient interactions revealed oligomers at high iSPOR concentrations, but it cannot be excluded that the crosslinked oligomers arose from random collisions (Supplementary Fig. 12). The published NMR structure of FtsN’s periplasmic domain appears to be monomeric^14^. Therefore, *in vitro* support for the SPOR-SPOR interaction we observed *in vivo* is lacking. One possible explanation is that the interaction requires one of the partners contain other domains of FtsN. Indeed, the Loose group reported that full-length FtsN *in vitro* forms a dimer in sedimentation experiments^40^, and the Vollmer group captured oligomeric FtsN molecules containing the periplasmic E and SPOR domains using blue native gel electrophoresis^51^.

### Cooperative binding of SPOR to dnG

Several findings support the idea that FtsN’s SPOR domain binds cooperatively to dnG. First, we observed that not only the stationary population percentages, but also the stationary lifetimes of single FtsN molecules, changed in response to a variety of altered conditions (Figs.1-3, Supplementary Fig. 13). The stationary lifetime, which reflects the time it takes for a single molecule to exit a particular state, directly measures the apparent dnG-dissociation time constant of FtsN. Changes in these lifetimes are therefore consistent with altered binding affinities between dnG and SPOR, in addition to shifted binding equilibria where only the bound and unbound population percentages change.

Second, although we could not directly quantify dnG level or length under different conditions, FtsN’s binding affinity responded to expected dnG levels in cells (Supplementary Fig. 13). For example, the stationary lifetime of FtsN increased to ∼60 s in the Δ6*LT*s mutant, where the dnG level is ∼20-40 fold higher than in WT cells (Fig. 1f, Supplementary Fig. 13)^19^. In the *ftsN*^ΔSPOR+^ background, iSPOR had a stationary lifetime similar to that of full-length FtsN (∼30 s), but in the presence of WT FtsN, iSPOR’s stationary lifetime decreased to ∼20 s and further to ∼14 s when cells were treated with aztreonam (Fig. 3d, bottom). In contrast, full-length FtsN’s stationary lifetime increased to ∼40 s in aztreonam-treated cells^23^. The decreased binding affinity of iSPOR to dnG in the presence of WT FtsN can be attributed to the expected low levels of dnG, as most dnG is occupied by full-length FtsN. The few-fold change in the binding affinities of SPOR to dnG, together with the *in vivo* evidences for SPOR-SPOR self-interaction, is most consistent with a cooperative binding mechanism, which has been frequently observed in DNA/mRNA-length-dependent cooperative binding of oligomeric proteins^41–43^. Notably, the Vollmer group previously demonstrated that the purified periplasmic domain of FtsN binds exclusively to dnG strands longer than 25 NAM-NAG units, consistent with a cooperative binding mechanism^18^.

Third, we considered alternative explanations for changes in the stationary lifetime of FtsN, specifically the possibility that these changes result from active “pulling” of SPOR by processive sPG synthases or “pushing” of SPOR by processive LTs, rather than cooperative binding of SPOR to dnG. We reasoned that active “pulling” by processive sPG synthases is unlikely because iSPOR (with FtsN^ΔSPOR+^), which lacks an E domain to bind to the sPG synthesis complex, had the same stationary lifetime as full-length WT FtsN (Fig. 3d, bottom). Similarly, active “pushing” of SPOR away from dnG by processive LTs is also unlikely since overexpression of iSPOR led to a strong chaining phenotype (Supplementary Fig. 8), suggesting that LTs are unable to “push” iSPOR off dnG.

Finally, all stationary lifetime distributions of FtsN under these conditions exhibited non-exponential distributions (Supplementary Fig. 10), indicating that the underlying process cannot be described as a single rate-limiting step. Many of these distributions showed a rise and decay, a signature indicative of multiple, consecutive steps^52^, which is consistent with a cooperative, non-monomeric process. These non-monotonic distributions suggest that the affinity of FtsN’s SPOR domain depends on the length of a dnG strand, and the long tails in the stationary lifetime plots (Supplementary Fig. 10) may arise from longer dnGs.

It remains unclear whether the self-interaction of SPOR on the sPG-track is also cooperative, but the persistent running time distributions of FtsN exhibited a rise and decay pattern similar to that of FtsN’s stationary lifetime distribution (Supplementary Fig. 3). Notably, FtsW’s slow-moving populations also showed non-exponential distributions (Supplementary Fig. 3). It is likely that these self-interactions are also cooperative, but further experiments are required to investigate this possibility.

### Biological function of cooperativity of SPOR

What would be the biological function of the cooperative self-interaction of SPOR domains? Cooperativity is often observed in biology systems when a switch-like behavior in response to a small change in protein or substrate concentration is required. For example, when LT activities are low (as mimicked by the Δ6*LT*s mutant), tight binding of FtsN to dnG can sequester nearly all sPG synthesis complexes on the dnG-track, thereby reducing the corresponding sPG synthesis activity. When sPG synthesis activity is low (as mimicked by aztreonam or fosfomycin treatment), the diminished slow-moving population of FtsN renders not only more FtsN molecules anchored to dnG but also tighter binding (stationary lifetime at ∼40 s), hence impeding LT activities on dnG (Fig. 4c, left). Conversely, when sPG synthesis activities are high (such as in the superfission mutant *ftsI*^R167S^, in the rich medium EZRDM, or under increased lipid II conditions), more FtsN molecules bind to the processively-moving sPG synthesis complex, making fewer FtsN molecules available to bind dnG and also lowering their binding affinities (stationary lifetime at ∼14 s). Consequently, more dnG is exposed to LTs for degradation, leading to higher degradation activities (Fig. 4c, right).

Furthermore, cooperativity can enhance the binding reaction of SPOR on either the dnG-track or the sPG-track, facilitating their respective functions. For example, truncation of SPOR led to a significant reduction in the slow-moving population of FtsW and the processivity of FtsN on the sPG-rack, suggesting that SPOR’s self-interaction on the sPG-track likely enhances the binding of FtsN to the synthesis complex.

We note that the truncation of SPOR did not impact the processivity of FtsW. The persistent running time of FtsW on the sPG-track was ∼20 s regardless of the presence or absence of FtsN’s SPOR domain (Supplementary Table 5). In contrast, the persistent running time of full-length FtsN was ∼14 s, significantly shorter than those of FtsW under the same conditions (Supplementary Tables 4 and 5). This discrepancy raises an interesting possibility, that is, multiple FtsN molecules may cycle within the processive sPG synthesis complex. While each FtsN molecule binds the complex for ∼14 s on average, there is always at least one FtsN molecule in the complex to maintain FtsWI’s processivity for ∼20 s. An example trajectory in Fig. 3k may indeed suggest this process. Furthermore, the slow-moving population percentage and persistent running time decreased for the dnG-binding defective FtsN^Q251E^-Halo^SW^ mutant and further for FtsN^ΔSPOR^-Halo^SW^ (Fig. 2), suggesting that SPOR’s binding to nearby dnG may facilitate the recycling process by restricting the diffusion of dissociated FtsN molecules.

Lastly, the cooperative binding of SPOR to dnG likely helps prevent the release of dissociated FtsWIQLB complex to the Z-track. However, the SPOR-dnG binding affinity needs to be in the right range. If the binding is too tight, as in the Δ6*LT*s background, all FtsWIQLB-FtsN complexes are sequestered on dnG, reducing sPG synthesis activity. Conversely, in the absence of SPOR-dnG binding, as in the Δ*amiABC* background, all complexes are sequestered on the Z-track, also diminishing sPG synthesis activity. We also note that as Δ6*LT*s and Δ*amiABC* cells may have other significant defects besides different dnG levels, further experiments are required to investigate these effects.

In summary, we provided strong evidence elucidating a mechanism for how FtsN may coordinate sPG synthesis and degradation activities in *E. coli*. FtsN controls the populations of sequestered and active sPG synthesis complexes on the dnG- and sPG-tracks by partitioning between the two tracks through the cooperative self-interaction of the SPOR domain. While other protein factors may still be at play, substrate availability and the natural binding affinities of FtsN to dnG and the synthesis complex can already provide a sensitive switch to regulate the partitioning of FtsN on the two tracks.

## Acknowledgements

We thank all members of the Xiao lab for helpful discussions and feedback on the manuscript, and members of the Weiss lab for help with strain construction. We would like to acknowledge use of resources at the Protein and Crystallography Facility within the Carver College of Medicine at the University of Iowa along with advice and assistance from Devin Reusch and Nicholas Schnicker. Work in the Xiao lab was supported by NIH R01GM086447 and R35GM136436 (to J.X.), GM125656 (subcontract to J.X.), a Hamilton Smith Innovative Research Award (to J.X.). Work in the Weiss lab was supported by NIH R01GM125656 (to D.S.W.).

## Author Contributions

Z.L., D.S.W. and J.X. conceived the study. Z.L., A.Y. and D.S.W. constructed the strains and performed genetic and phenotypic experiments. Z.L. and X.Y. performed the imaging experiments and analyzed the data. S.H. performed the HADA labeling and imaging experiments. J.W.M. wrote the custom MATLAB script for speed deconvolution in analyzing the single-molecule tracking data. X.C. performed the crosslinking experiments. X.Y. provided the single-molecule tracking data of Halo-FtsW in *C. crescentus*. B. H. B. provided the single-molecule tracking data of Halo-FtsB in *E. coli*. Z.L., D.S.W. and J.X. wrote the original draft. All authors reviewed and edited the manuscript. D.S.W. and J.X. supervised the study. Funding was acquired by D.S.W. and J.X.

## References

1. Vollmer, W. & Bertsche, U. Murein (peptidoglycan) structure, architecture and biosynthesis in *Escherichia coli*. Biochim. Biophys. Acta 1778, 1714–1734 (2008).

2. van Heijenoort, J. Peptidoglycan hydrolases of *Escherichia coli*. Microbiol. Mol. Biol. Rev. 75, 636–663 (2011).

3. Jorgenson, M.A., Chen, Y., Yahashiri, A., Popham, D.L. & Weiss, D.S. The bacterial septal ring protein RlpA is a lytic transglycosylase that contributes to rod shape and daughter cell separation in *Pseudomonas aeruginosa*. Mol. Microbiol. 93, 113–128 (2014).

4. Heidrich, C. et al. Involvement of N-acetylmuramyl-L-alanine amidases in cell separation and antibiotic-induced autolysis of *Escherichia coli*. Mol. Microbiol. 41, 167–178 (2001).

5. Priyadarshini, R., de Pedro, M.A. & Young, K.D. Role of peptidoglycan amidases in the development and morphology of the division septum in *Escherichia coli*. J. Bacteriol. 189, 5334–5347 (2007).

6. Scheurwater, E., Reid, C.W. & Clarke, A.J. Lytic transglycosylases: bacterial space-making autolysins. Int. J. Biochem. Cell Biol. 40, 586–591 (2008).

7. Vollmer, W., Joris, B., Charlier, P. & Foster, S. Bacterial peptidoglycan (murein) hydrolases. FEMS Microbiol. Rev. 32, 259–286 (2008).

8. Busiek, K.K. & Margolin, W. A role for FtsA in SPOR-independent localization of the essential *Escherichia coli* cell division protein FtsN. Mol. Microbiol. 92, 1212–1226 (2014).

9. Liu, B., Persons, L., Lee, L. & de Boer, P.A. Roles for both FtsA and the FtsBLQ subcomplex in FtsN-stimulated cell constriction in *Escherichia coli*. Mol. Microbiol. 95, 945–970 (2015).

10. Pichoff, S., Du, S. & Lutkenhaus, J. The bypass of ZipA by overexpression of FtsN requires a previously unknown conserved FtsN motif essential for FtsA-FtsN interaction supporting a model in which FtsA monomers recruit late cell division proteins to the Z ring. Mol. Microbiol. 95, 971–987 (2015).

11. Dai, K., Xu, Y. & Lutkenhaus, J. Topological characterization of the essential *Escherichia coli* cell division protein FtsN. J. Bacteriol. 178, 1328–1334 (1996).

12. Gerding, M.A. et al. Self-enhanced accumulation of FtsN at division sites and roles for other proteins with a SPOR domain (DamX, DedD, and RlpA) in Escherichia coli cell constriction. J. Bacteriol. 191, 7383–7401 (2009).

13. Park, K.T., Johnson, D.K., Pichoff, S., Du, S. & Lutkenhaus, J. The essential domain of FtsN triggers cell division by promoting interaction between FtsL and FtsI. bioRxiv (2023).

14. Yang, J.C., Van Den Ent, F., Neuhaus, D., Brevier, J. & Lowe, J. Solution structure and domain architecture of the divisome protein FtsN. Mol. Microbiol. 52, 651–660 (2004).

15. Britton, B.M. et al. Conformational changes in the essential *E. coli* septal cell wall synthesis complex suggest an activation mechanism. Nat. Commun. 14, 4585–4599 (2023).

16. Alcorlo, M. et al. Structural basis of denuded glycan recognition by SPOR domains in bacterial cell division. Nat. Commun. 10, 5567–5579 (2019).

17. Moll, A. & Thanbichler, M. FtsN-like proteins are conserved components of the cell division machinery in proteobacteria. Mol. Microbiol. 72, 1037–1053 (2009).

18. Ursinus, A. et al. Murein (peptidoglycan) binding property of the essential cell division protein FtsN from *Escherichia coli*. J. Bacteriol. 186, 6728–6737 (2004).

19. Yahashiri, A., Jorgenson, M.A. & Weiss, D.S. Bacterial SPOR domains are recruited to septal peptidoglycan by binding to glycan strands that lack stem peptides. Proc. Natl Acad. Sci. USA 112, 11347–11352 (2015).

20. McCausland, J.W. et al. Treadmilling FtsZ polymers drive the directional movement of sPG-synthesis enzymes via a Brownian ratchet mechanism. Nat. Commun. 12, 609–621 (2021).

21. Yang, X. et al. A two-track model for the spatiotemporal coordination of bacterial septal cell wall synthesis revealed by single-molecule imaging of FtsW. Nat. Microbiol. 6, 584–593 (2021).

22. Yang, X. et al. GTPase activity-coupled treadmilling of the bacterial tubulin FtsZ organizes septal cell wall synthesis. Science 355, 744–747 (2017).

23. Lyu, Z. et al. FtsN maintains active septal cell wall synthesis by forming a processive complex with the septum-specific peptidoglycan synthases in *E. coli*. Nat. Commun. 13, 5751–5766 (2022).

24. Mahone, C.R. et al. Integration of cell wall synthesis and chromosome segregation during cell division in *Caulobacter*. J. Cell Biol. 223, e20221 (2024).

25. Perez, A.J. et al. Movement dynamics of divisome proteins and PBP2x:FtsW in cells of *Streptococcus pneumoniae*. Proc. Natl Acad. Sci. USA 116, 3211–3220 (2019).

26. Whitley, K.D. et al. Peptidoglycan synthesis drives a single population of septal cell wall synthases during division in *Bacillus subtilis*. Nat. Microbiol. 9, 1064–1074 (2024).

27. Schaper, S. et al. Cell constriction requires processive septal peptidoglycan synthase movement independent of FtsZ treadmilling in *Staphylococcus aureus*. Nat. Microbiol. 9, 1049–1063 (2024).

28. Perez, A.J. & Xiao, J. Stay on track - revelations of bacterial cell wall synthesis enzymes and things that go by single-molecule imaging. Curr. Opin. Microbiol. 79, e102490 (2024).

29. Park, K.T., Du, S. & Lutkenhaus, J. Essential role for FtsL in activation of septal peptidoglycan synthesis. mBio 11, e03012 (2020).

30. Condon, S.G.F. et al. The FtsLB subcomplex of the bacterial divisome is a tetramer with an uninterrupted FtsL helix linking the transmembrane and periplasmic regions. J. Biol. Chem. 293, 1623–1641 (2018).

31. Marmont, L.S. & Bernhardt, T.G. A conserved subcomplex within the bacterial cytokinetic ring activates cell wall synthesis by the FtsW-FtsI synthase. Proc. Natl Acad. Sci. USA 117, 23879–23885 (2020).

32. Navarro, P.P. et al. Cell wall synthesis and remodelling dynamics determine division site architecture and cell shape in *Escherichia coli*. Nat. Microbiol. 7, 1621–1634 (2022).

33. Yahashiri, A., Kaus, G.M., Popham, D.L., Houtman, J.C.D. & Weiss, D.S. Comparative study of bacterial SPOR domains identifies functionally important differences in glycan binding affinity. J. Bacteriol. 204, e0025222 (2022).

34. Peters, N.T., Dinh, T. & Bernhardt, T.G. A fail-safe mechanism in the septal ring assembly pathway generated by the sequential recruitment of cell separation amidases and their activators. J. Bacteriol. 193, 4973–4983 (2011).

35. Yakhnina, A.A. & Bernhardt, T.G. The Tol-Pal system is required for peptidoglycan-cleaving enzymes to complete bacterial cell division. Proc. Natl Acad. Sci. USA 117, 6777–6783 (2020).

36. Yahashiri, A., Jorgenson, M.A. & Weiss, D.S. The SPOR domain, a widely conserved peptidoglycan binding domain that targets proteins to the site of cell division. J. Bacteriol. 199, e00118 (2017).

37. Squyres, G.R. et al. Single-molecule imaging reveals that Z-ring condensation is essential for cell division in *Bacillus subtilis*. Nat. Microbiol. 6, 553–562 (2021).

38. Buss, J. et al. A multi-layered protein network stabilizes the *Escherichia coli* FtsZ-ring and modulates constriction dynamics. PLoS Genetics 11, e1005128 (2015).

39. Barrows, J.M., Anderson, A.S., Talavera-Figueroa, B.K. & Goley, E.D. Intrinsic and extrinsic factors regulate FtsZ function in Caulobacter crescentus. bioRxiv (2023).

40. Baranova, N. et al. Diffusion and capture permits dynamic coupling between treadmilling FtsZ filaments and cell division proteins. Nat. Microbiol. 5, 407–417 (2020).

41. Peisley, A. et al. Kinetic mechanism for viral dsRNA length discrimination by MDA5 filaments. Proc. Natl Acad. Sci. USA 109, 3340–3349 (2012).

42. Morrone, S.R. et al. Cooperative assembly of IFI16 filaments on dsDNA provides insights into host defense strategy. Proc. Natl Acad. Sci. USA 111, 62–71 (2014).

43. Hooy, R.M. & Sohn, J. The allosteric activation of cGAS underpins its dynamic signaling landscape. eLife 7, e39984 (2018).

44. Duncan, T.R., Yahashiri, A., Arends, S.J., Popham, D.L. & Weiss, D.S. Identification of SPOR domain amino acids important for septal localization, peptidoglycan binding, and a disulfide bond in the cell division protein FtsN. J. Bacteriol. 195, 5308–5315 (2013).

45. Du, S., Pichoff, S. & Lutkenhaus, J. FtsEX acts on FtsA to regulate divisome assembly and activity. Proc. Natl Acad. Sci. USA 113, 5052–5061 (2016).

46. Mannik, J., Pichoff, S., Lutkenhaus, J. & Mannik, J. Cell cycle-dependent recruitment of FtsN to the divisome in *Escherichia coli*. mBio 13, e0201722 (2022).

47. Kashammer, L. et al. Cryo-EM structure of the bacterial divisome core complex and antibiotic target FtsWIQBL. Nat. Microbiol. 8, 1149–1159 (2023).

48. Nguyen, H.T.V. et al. Structure of the heterotrimeric membrane protein complex FtsB-FtsL-FtsQ of the bacterial divisome. Nat. Commun. 14, 1903–1915 (2023).

49. Li, G.W., Burkhardt, D., Gross, C. & Weissman, J.S. Quantifying absolute protein synthesis rates reveals principles underlying allocation of cellular resources. Cell 157, 624–635 (2014).

50. Arends, S.J. et al. Discovery and characterization of three new *Escherichia coli* septal ring proteins that contain a SPOR domain: DamX, DedD, and RlpA. J. Bacteriol. 192, 242–255 (2010).

51. Muller, P. et al. The essential cell division protein FtsN interacts with the murein (peptidoglycan) synthase PBP1B in *Escherichia coli*. J. Biol. Chem. 282, 36394–36402 (2007).

52. Floyd, D.L., Harrison, S.C. & van Oijen, A.M. Analysis of kinetic intermediates in single-particle dwell-time distributions. Biophys. J. 99, 360–366 (2010).

