## Supplementary Information for "*E. coli* FtsN coordinates synthesis and degradation of septal peptidoglycan by partitioning between a synthesis track and a denuded glycan track"

### Methods

**Growth media.** Lysogeny broth (LB) was employed for routine growth and genetic manipulation of *E. coli* strains. For microscopy, cells were grown in the M9-glucose minimal medium (0.4% D-glucose, 1× MEM amino acids and 1× MEM vitamin)[1]. For single-molecule tracking experiments, vitamins were omitted from the M9 medium (termed $M9^{-}$) to minimize background fluorescence. Where appropriate, antibiotics were included as follows: ampicillin, 100 μg $mL^{-1}$; carbenicillin, 25 μg $mL^{-1}$; chloramphenicol, 35 μg $mL^{-1}$; kanamycin, 40 μg $mL^{-1}$; spectinomycin, 100 μg $mL^{-1}$ for plasmids and 35 μg $mL^{-1}$ for chromosomal alleles. In general antibiotics were used when propagating *E. coli* strains in LB but omitted when growing cells in minimal media for microscopy. This was possible because most of the fusions used in this study were integrated into the chromosome. However, chloramphenicol was added to minimal media for growth of strains with plasmids pJL130, pJL132, pJL137, or pJL139. Where appropriate, 0.2% L-arabinose was used to express *ftsN* under $P_{BAD}$ control[2]. Isopropyl β-D-1-thiogalactopyranoside (IPTG) was used as indicated to express genes under control of modified (weakened) *Trc* promoters.

**Bacterial strains, oligonucleotide primers, and plasmids.** Standard procedures were used for analysis of DNA, PCR, electroporation, transformation, P1 transduction and integration of CRIM plasmids[3]. Bacterial strains are listed in Supplementary Table 1, which also describes strain construction. Plasmids are described in Supplementary Table 2, followed by descriptions of how these plasmids were made. Some plasmids were assembled by amplifying appropriate DNA fragments using Q5 DNA polymerase followed by ligation or assembly into restriction-digested vectors using T4 DNA ligase or NEBuilder HiFi DNA Assembly Master Mix, respectively (New England Biolabs). Alternatively, plasmids were assembled by amplifying appropriate insert and vector DNA fragments followed by In-Fusion cloning (Takara, In-Fusion HD Cloning Kit). The QuikChange Lightening Kit (Aligent) was used for site-directed mutagenesis as needed. Oligonucleotides were from Integrated DNA Technologies (Coralville, IA), and are listed in Supplementary Table 3. Regions of plasmids encompassing fusion genes constructed by PCR were verified by DNA sequencing.

**Growth of cells for microscopy.** Starter cultures were grown overnight at 30 °C in LB. Where appropriate, LB was supplemented with antibiotics and 0.2% L-arabinose to induce

$P_{BAD}$::*ftsN*. The next day cells were washed once with M9-glucose to remove antibiotics and arabinose, then diluted 500 to 2000-fold into 3 mL of M9-glucose supplemented with 10 μM IPTG to induce the expression of chromosomal *ftsN* fusions, or without IPTG to permit the leaky expression of *ftsN* and *ftsW* fusions from the plasmid; antibiotics were excluded, except for strains harboring plasmids. Cultures were incubated at room temperature (RT) with shaking until the $OD_{600}$ reached ~0.35 (~18 hours).

**Growth curve measurements.** Starter cultures were grown overnight at 30 °C in M9-glucose. The following day, $OD_{600}$ was measured with a Nanodrop and cultures were diluted to an initial $OD_{600}$ of 0.1 in 200 µl M9-glucose in a Corning Costar sterile 96-well plate. The 96-well plates were incubated and shaken in a Tecan Infinite M200 Pro set at 30°C, with $OD_{600}$ measurements taken every 30 min for 23.5 h, shaking the plate for 3 min at 220 rpm before measuring.

**HADA labeling and imaging.** Starter cultures were grown overnight at 30 °C in LB, with kanamycin being added only to *ftsN*$^{\Delta SPOR+}$ cells. The next day cells were washed once with M9-glucose and diluted 1:200 for *ftsN*$^{\Delta SPOR}$, Δ*amiABC*, and Δ6*LT*s strains, and 1:500 for WT and *ftsN*$^{\Delta SPOR+}$ strains into 3 mL of M9-glucose. Cultures were incubated at RT with shaking until the $OD_{600}$ reached ~0.35.

Cultures under the cephalexin condition were then treated with cephalexin (14 mg $mL^{-1}$) for a final concentration of 20 µg $mL^{-1}$. These cultures were incubated at RT for 2 hours with shaking at 240 rpm. Cells with and without cephalexin treatment were then incubated with 80 µM HADA for 5 minutes on a covered nutator at 25 °C before being fixed with 70% ethanol. The cells were incubated for 20 minutes on a covered nutator at 4 °C before being washed 3 times and resuspended in 1x PBS. Previous experiments with HADA revealed 5-minutes labeling time to be the most viable option (as opposed to 2- or 10-minutes labeling times) as it prevented sick strains such as Δ*amiABC* and Δ6*LT*s from oversaturating the camera while also avoiding a loss of septum labeling.

3% PBS gels were prepared by heating a mixture of SeaPlaque GTG Agarose powder and 1x PBS to 70 °C for 30 minutes in a heat block. 75 µL of the gel were loaded onto an acrylic coverslip encased in a plastic chamber. A rubber gasket and glass coverslip were placed around and on top of the gel, respectively, for 30 minutes. The gel was then cut into 8 uniform slices and the samples were loaded. After 3 minutes, a new glass coverslip was applied, and the construct was encased in a metal chamber. The sample was then kept in the microscope room for 30 minutes to reach equilibrium with its environment.

The Olympus IX-71 scope, 1.30 NA oil Ph3 100x phase objective, 405-nm laser (20 W $cm^2$), and Andor EmCCD camera (16 µm $pixel^{-1}$) were used to image the strains. No emission filter was used. For each strain, 5-15 regions were imaged using a program which captured brightfield, fluorescent, and post-brightfield images.

The percentage of septal HADA insertion was assessed by first measuring the integrated fluorescent intensities of the septum ($I_{septum}$) and the whole cell ($I_{cell}$) after subtracting the background fluorescent signal close to the cells. Since Δ*amiABC* and Δ6*LT*s chaining cells have multiple septa, the density of septa in individual cells obtained through dividing the septa number by the cell length was used to calibrate the difference of septum numbers. A scaling factor $\alpha$ was generated by normalizing the density of septa to that of the WT cells. The percentage of septal HADA insertion was then calculated as $(I_{septum})/(I_{cell})/\alpha$.

**Cell-labeling with Janelia Fluor 646 (JF646) dye**. Cells from 1 mL of culture at an $OD_{600}$ ~0.35 were harvested by centrifugation and the cell pellet was resuspended with 1 mL $M9^{-}$ minimal medium including 1 nM JF646 (for SMT imaging) or 1 µM JF646 (for ensemble imaging). The culture was mixed well with a pipette and put on a nutator at RT for 30 minutes. After labeling, cells were washed three times with $M9^{-}$ minimal medium and concentrated to a final volume of 50 µL after resuspending the cell pellets in fresh with $M9^{-}$ minimal medium.

**Time-lapse imaging.** Cells were grown and labeled as described above. 0.5 µL of 100 µg $mL^{-1}$ aztreonam solution was added to the 50 µL concentrated labeled cells. The final concentration was 1 µg $mL^{-1}$. 0.5 µL of 100 µg $mL^{-1}$ aztreonam solution was also added on top of a 50-µL, 3% agarose gel pad right before applying cells. The moment of the latter was counted as time zero. Time-lapse imaging was performed on a home-built microscope as previously described[4]. Images were acquired every 1-minute for 6 hours, with a 100-ms exposure time.

**SMT imaging and data analysis**. Cells were grown and labeled as described above except for certain conditions listed below. Cells were then loaded onto a 50-µL, 3% agarose gel pad (containing the same growth medium without antibiotic) laid in an observation chamber (FCS2, Bioptechs). The chamber was locked on the microscope stage (ASI, Eugene, OR) to minimize mechanical drift.

For aztreonam-treated conditions, 0.5 μL of 100 μg $mL^{-1}$ aztreonam solution was added to the 50 μL concentrated labeled cells. The final concentration was 1 μg $mL^{-1}$. 0.5 μL of 100 μg $mL^{-1}$ aztreonam solution was also added on top of the gel pad right before applying cells. The moment of the latter was counted as time zero. The chamber with cells was kept on the microscope stage for 30 minutes before the images were acquired. All images were collected within 3 hours of drug treatment.

SMT imaging was performed in the same way as previously described[4]. Briefly, the tracking imaging was performed on an Olympus IX71 inverted microscope equipped with a 100x, 1.49 NA oil-immersion objective and Andor iXon 897 Ultra EM-CCD camera in epifluorescence-illumination mode using Metamorph 7.8.13.0 software. The focal plane was placed at ~250 nm from the bottom of the cell to image the molecules moving on the bottom half of the cylindrical portion of the cell body. Single molecules were tracked with 100-ms exposure time using a 647-nm laser with intensity 30 W $cm^{-2}$. 240 frames were acquired with 1Hz frame rate. 3D imaging was achieved as described previously[4, 5].

The data processing was similar as previously described[4, 6]. Specifically, the xy positions of single molecules were determined through the 2D Gaussian fitting of the Point Spread Function (PSF) with ThunderSTORM[7], a plug-in for ImageJ[8], while the z position was given by the calibration curve obtained by z-scanning of the fluorescent microspheres[9]. Because of the refractive index mismatch between the transmission path of the microspheres used for calibration (glass and oil) and that of the fluorescent proteins (aqueous cell environment, glass, and oil), the z values obtained from the calibration curve were rescaled by a factor of 0.75. A bandpass filter (60-300 nm) for both sigma1 and sigma2 was applied to remove the single pixel noise and out-of-focus molecules. All analysis thereafter used custom scripts in MATLAB R2020a. The localizations were linked to trajectories using a home-built MATLAB script (https://zenodo.org/records/8189840) that adopted the nearest neighbor algorithm[10]. The distance threshold was set to 300 nm per frame, which approximates to a max diffusion coefficient of ~0.05 $\mu m^2 s^{-1}$, or a max speed of 300 nm $s^{-1}$. To link molecules which may have blinked across frames or left the focal plane, a time threshold of 8 frames was chosen according to the off-time distribution. Only trajectories near the midpoint of the cell's long axis or near visible constriction sites where cell division takes place were used in the analysis to ensure the molecules are cell division and septal peptidoglycan related.

Due to the rod-shape cell envelope, the real displacement of the tracked molecules around the circumference is underestimated. The trajectories were unwrapped to one dimension using a home-built MATLAB script (https://zenodo.org/records/8189840). We noticed that the velocity estimated from MSD curve fitting is not accurate when the dwell time of the trajectory is short (< 20 frames) or when there is more than one moving state in a single trajectory. Unwrapped trajectories were then segmented manually to determine whether a single molecule in a segment is stationary or moving processively as previously described[6]. Briefly, a segment was first fitted with a line. R, which is the ratio of the displacement and the standard deviation of fitting residuals, and P, which is the probability of processive movement, were defined and used as the criteria for the classification of segments. Through manual inspection, we determined to classify segments as processive based on a threshold of R ≤ 0.4 and P ≥ 0.5, while all others were classified as stationary. Since the confidence of classification is correlated with the segmentation length, we only consider segments longer than 5 frames to minimize classification error.

The cumulative probability density (CDF) of directional moving FtsN speeds were calculated for each condition and fit to either a single or double log-normal populations:

$$CDF = P_1 \frac{\left(1 + \mathrm{erf}\left[\frac{ln\, v - u_1}{\sqrt{2\sigma_1}}\right]\right)}{2} + (1 - P_1)\frac{\left(1 + \mathrm{erf}\left[\frac{ln\, v - u_2}{\sqrt{2\sigma_2}}\right]\right)}{2}$$

Where $v$ is the moving speed for FtsN or it's mutants and $P_1$ is the percentage of the first population. For fitting with a single population, $P_1 = 1$. The values $u$ and $\sigma$ are the natural logarithmic mean and standard deviation. The average speed of each population is calculated as $\exp\left(u + \frac{\sigma^2}{2}\right)$. To estimate the error of the speed and percentage (Supplementary Tables 4-6), the CDF curves were bootstrapped 200 times and fit with the corresponding equation (single- or double-population).

To fit the stationary population, histograms were generated of the velocities from corresponding stationary segments in the respective FtsN condition data. Peak values and bin centers were used to fit a single exponential decay function:

$$f(x) = A * \exp\,(-\lambda x)$$

Where A is the amplitude of the fitted curve and $\lambda = \frac{1}{u}$. $u$ is the mean "velocity" of stationary segments.

Deconvolution of the fast- and slow-moving populations (Fig. 2c) was performed in the same way as previously described[11].

**Calculation of FtsN's occupancy on the sPG synthesis track and dnG.** We calculated FtsN's occupancy on the processive sPG-track ($N_{\mathrm{sPG}}$) using the following equation:

$$N_{\mathrm{sPG}} = P_{\mathrm{sm}} \cdot T_{\mathrm{sm}} \cdot V_{\mathrm{sm}}$$

Where $P_{\mathrm{sm}}$ is the percentage of the slow-moving population of FtsN, $T_{\mathrm{sm}}$ is the processive rum time (s) of the slow-moving population, and $V_{\mathrm{sm}}$ is the velocity (nm $s^{-1}$) of the slow-moving population. The $N_{\mathrm{sPG}}$value of each condition was then normalized to that of the WT condition (FtsN-Halo$^{SW}$ in MG1655, strain EC5234, Supplementary Table 1). Errors in the final normalized $N_{\mathrm{sPG}}$ values were propagated from experimentally determined errors in each parameter. FtsW's occupancy on the processive sPG-track ($W_{\mathrm{sPG}}$) was calculated in the same way.

We used FtsN's occupancy on the processive sPG-track as a proxy for sPG synthesis activity. Previously, we showed that FtsW's occupancy on the sPG-track correlated to the cell wall constriction rate and HADA incorporation level[6]. Here, we also showed that FtsN's occupancy on the sPG-track correlated with FtsW's occupancy on the sPG-track (Supplementary Fig. 9).

Similarly, we calculated FtsN's occupancy on dnG using the following equation:

$$N_{\mathrm{dnG}} = P_{\mathrm{stat}} \cdot T_{\mathrm{stat}}$$

Where $P_{\mathrm{stat}}$ is the percentage of the stationary, dnG-bound population of FtsN and $T_{\mathrm{stat}}$ is the corresponding stationary life time (s). The $N_{\mathrm{dnG}}$value of each condition was then normalized to that of the WT condition (FtsN-Halo$^{SW}$ in MG1655, strain EC5234, Supplementary Table 1). Errors in the final normalized $N_{\mathrm{dnG}}$ values were propagated from experimentally determined errors in each parameter. This calculation only applies to FtsN constructs that have an intact SPOR domain.

The $N_{\mathrm{dnG}}$value reflects the existing level of dnG available for FtsN to bind. The higher the $N_{\mathrm{dnG}}$value, the higher the level of dnG, and hence the lower activities of LTs, which degrade dnG. Therefore, we used $1/N_{\mathrm{dnG}}$ as a proxy for LT's sPG degradation activity.

**Purification of FtsN$^{SPOR}$ domain protein (iSPOR).** Wild-type $His_6$-tagged iSPOR protein was overproduced in *E. coli* SHuffle T7 (New England BioLabs) and purified at 4 °C by cobalt affinity chromatography as described[12]. Purified protein was dialyzed into 25 mM $Na_2PO_4$, 150 mM NaCl, pH 7.5 buffer. Aliquots were analyzed by size exclusion chromatography and static light scattering immediately without freezing.

**Size Exclusion Chromatography.** Size exclusion chromatography was performed using a Cytiva Superdex™ 75 10/300GL 24-mL column. The column was equilibrated with 30 mL of buffer containing 25 mM $Na_2PO_4$, 150 mM NaCl, pH 7.5 buffer. Once equilibrated a volume of 0.5 mL of iSPOR was loaded on the column using a 0.5-mL loop. The column was run at 4 °C at a rate of 0.5 mL $min^{-1}$ for 30 mL. The approximate molecular weight was determined from Bio-Rad Gel Filtration Standard (Cat. #1511901).

**Static Light Scattering.** Static light scattering was performed using a DynaPro® NanoStar® (Wyatt) instrument in 25 mM $Na_2PO_4$, 150 mM NaCl, pH 7.5 buffer. The iSPOR samples were prepared by concentrating the size exclusion chromatography peak to 1.7 mg $mL^{-1}$. The protein was loaded into a quartz cuvette at a volume of 10 µL. The cuvette was placed in the instrument and the temperature was stabilized to 20°C. After stabilization, 10 measurements were taken to get the average number.

**iSPOR labeling with Cy3B.** Labelling of iSPOR with Cy3B dye was performed by incubating 100 µM iSPOR with 3-fold molar excess of mono-reactive NHS ester-modified Cy3B (PA63101, Cytiva) in 100 mM $NaHCO_3/Na_2CO_3$, pH 8.3 buffer in the dark at 25 °C for 2 hours. Excess dye was removed using a PD-10 desalting column (17085101, Cytiva) in 25 mM $Na_2PO_4$, 150 mM NaCl, pH 7.5 buffer. The dye-to-protein molar ratio of the eluted protein sample was determined by dividing the molar concentration of Cy3B by that of iSPOR after correction according to the product instruction. The labelling ratio is 1.1 for iSPOR-Cy3B.

**iSPOR / iSPOR-Cy3B cross-linking with glutaraldehyde.** 300 µM of iSPOR protein was diluted to final concentrations of 100 µM, 30 µM, 10 µM, 3 µM, 1 µM, 300 nM, 100 nM, 30 nM, 10 nM, 3 nM, and 1 nM. Glutaraldehyde was added into 50 µL of above protein samples to a final concentration of 0.1% (v/v). The reactions were incubated in the dark at 25 °C for 15 minutes. Samples were mixed with protein loading buffer and heated for 10 min at 95 °C before loading 10 µl onto a precast mini-PROTEAN TGX gel (10% polyacrylamide, from Bio-Rad, Hercules, CA). The protein gel was stained with Coomassie Brilliant blue R 250 (Supplementary Figure 12a).

Since the protein detection limit using Coomassie staining is too low, iSPOR-Cy3B sample was used as described below. 10 µM of iSPOR-Cy3B sample was diluted to final concentrations of 3 µM, 1 µM, 300 nM, 100 nM, 30 nM, 10 nM, 3 nM, and 1 nM.

Glutaraldehyde was added into 50 µL of above protein samples to a final concentration of 0.1% (v/v). The reactions were incubated in the dark at 25 °C for 15 minutes. Samples were mixed with protein loading buffer and heated for 10 min at 95 °C before loading 10 µl onto a precast mini-PROTEAN TGX gel (10% polyacrylamide, from Bio-Rad, Hercules, CA). The protein gel was imaged on a Typhoon biomolecular imager directly without Coomassie staining (Supplementary Figure 12b).

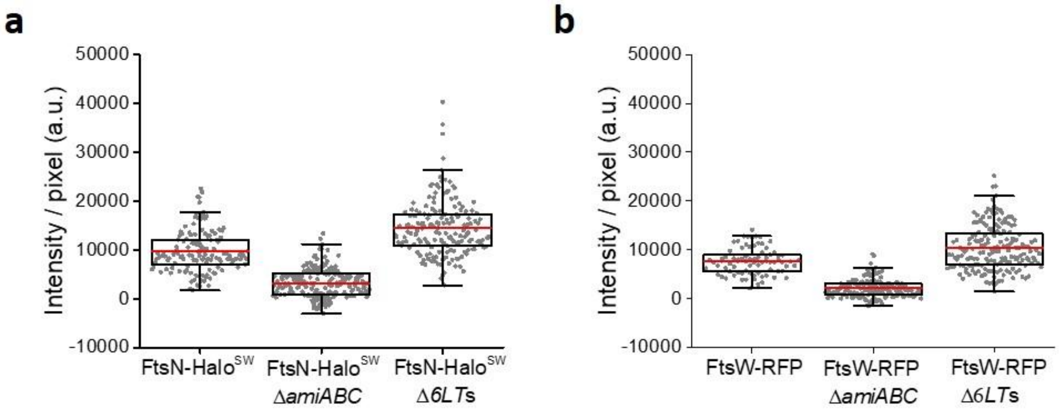


**Supplementary Fig. 1. Pixel intensity of FtsN-Halo$^{SW}$ and FtsW-RFP in WT, Δ*amiABC*, and Δ6*LT*s strains.**

The pixel intensity of FtsN-Halo$^{SW}$ (**a**) and FtsW-RFP (**b**) in WT, Δ*amiABC*, and Δ6*LT*s strains after subtracting the background fluorescence in cytoplasm.

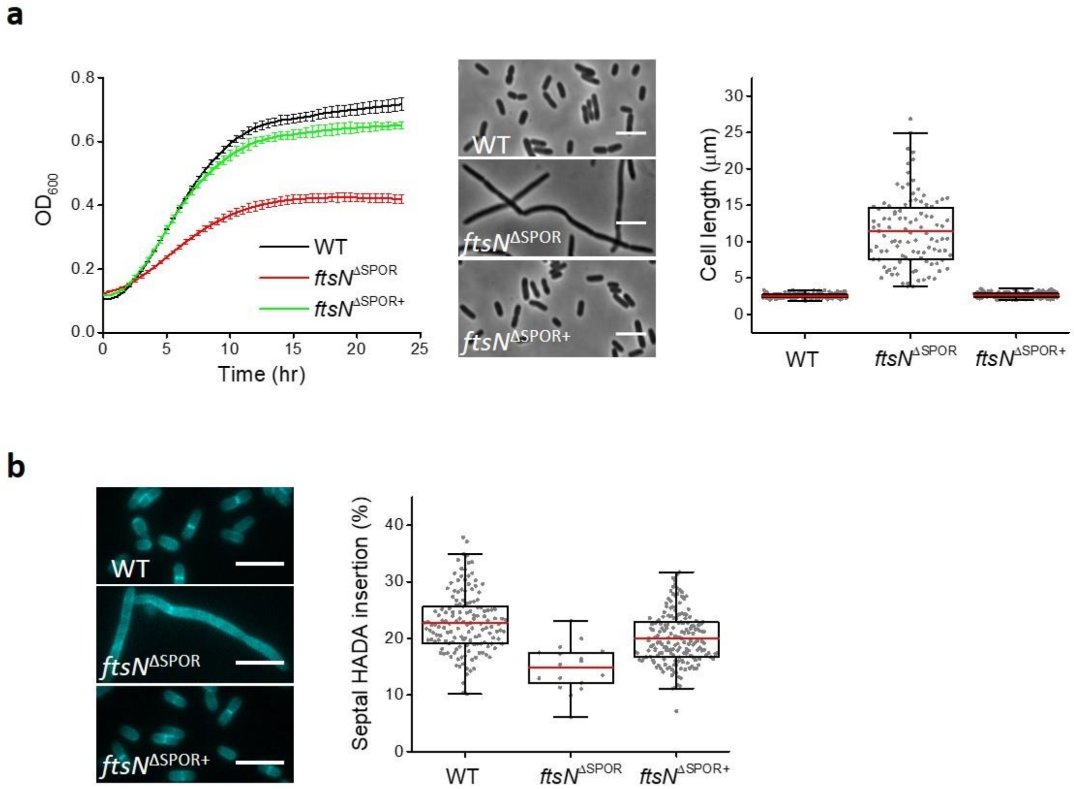


**Supplementary Fig. 2. Extra copies of FtsN$^{\Delta SPOR}$ restored cell growth and sPG synthesis.**

(**a**) Cell growth curves, phase contrast images, and cell length of WT, *ftsN*$^{\Delta SPOR}$, and *ftsN*$^{\Delta SPOR+}$ strains. (**b**) Representative HADA labeling images of WT, *ftsN*$^{\Delta SPOR}$, and *ftsN*$^{\Delta SPOR+}$ cells and the quantification of HADA insertion in septum compared to the whole cell. Scale bars, 5 µm.

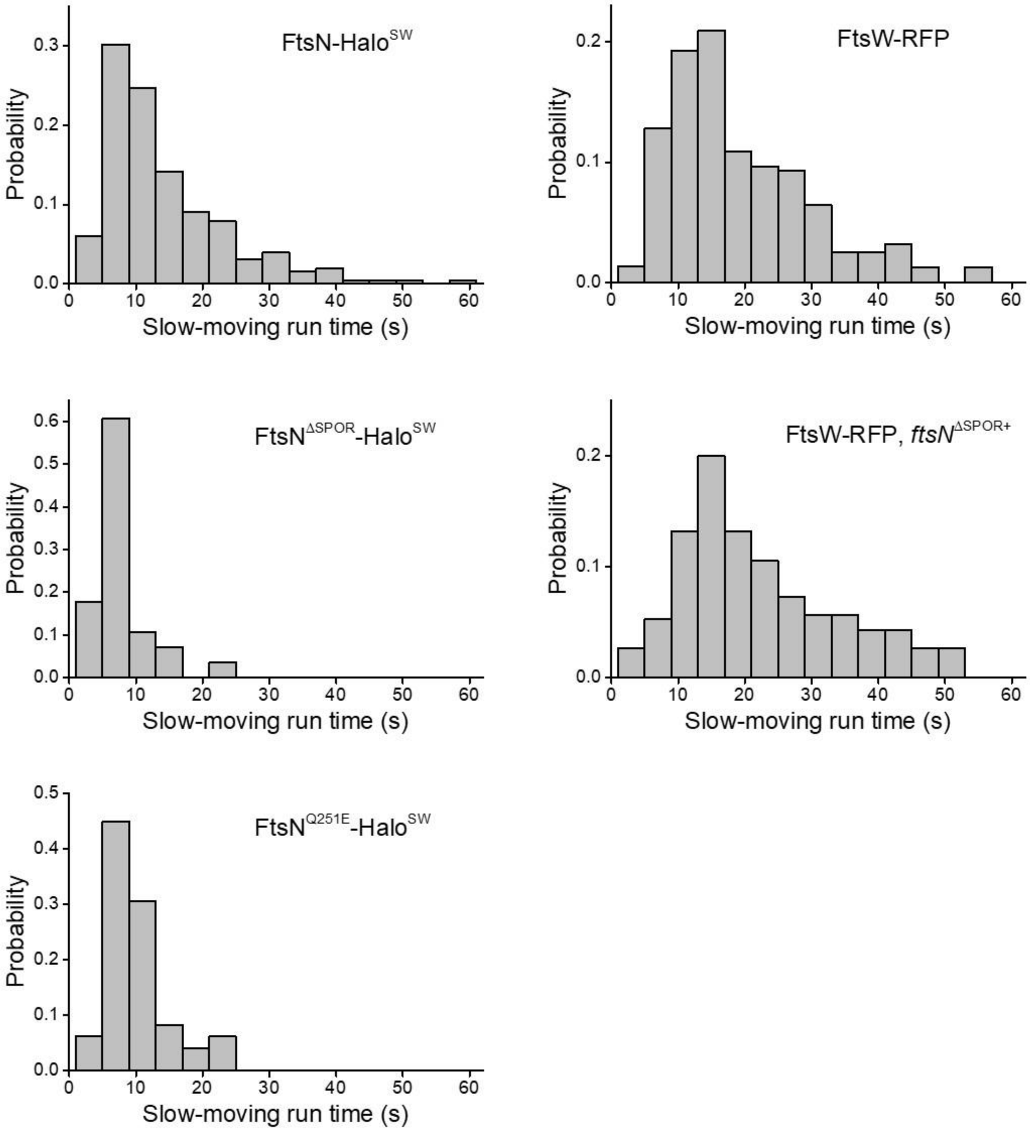


**Supplementary Fig. 3. The persistent running time distributions of the slow-moving population of FtsW and FtsN variants under different conditions.**

The persistent running time distributions of FtsW and FtsN variants under various conditions exhibited non-exponential distributions.

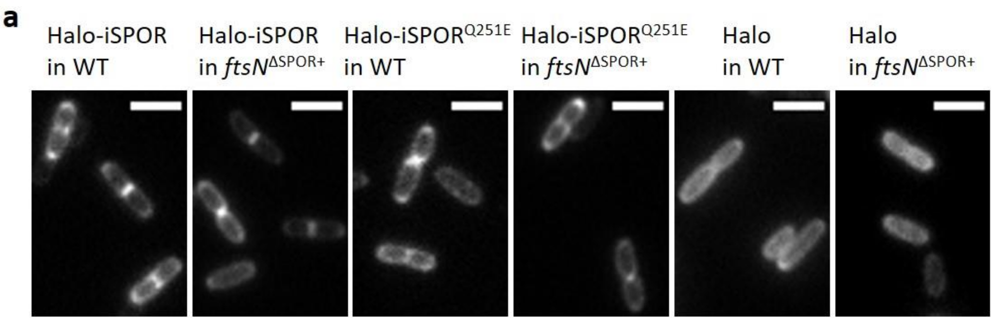


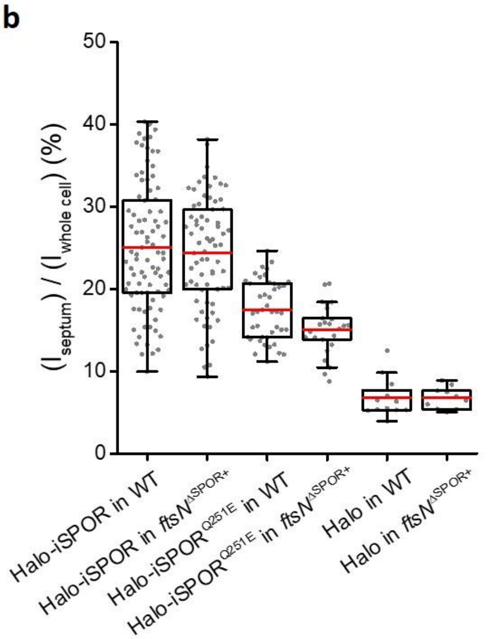


| Constructs | Cells with visible iSPOR localization ($N_{iSPOR}/N_{total}$) | $I_{septum}$ / $I_{whole\ cell}$ |
|---|---|---|
| Halo-iSPOR in WT | 43.2 % (89/206) | 25.0 ± 0.8 % |
| Halo-iSPOR in *ftsN*$^{\Delta SPOR+}$ | 43.7 % (73/167) | 24.4 ± 0.8 % |
| Halo-iSPOR$^{Q251E}$ in WT | 23.1 % (43/186) | 17.4 ± 0.5 % |
| Halo-iSPOR$^{Q251E}$ in *ftsN*$^{\Delta SPOR+}$ | 20.6% (21/102) | 15.1 ± 0.6 % |
| Halo in WT | 4.8 % (12/249) | 6.8 ± 0.1 % |
| Halo in *ftsN*$^{\Delta SPOR+}$ | 5.2 % (10/192) | 6.8 ± 0.1 % |

**Supplementary Fig. 4. Septal localization of Halo-iSPOR and Halo-iSPOR$^{Q251E}$ in WT and *ftsN*$^{\Delta SPOR+}$ strains.**

(**a**) Representative ensemble images of Halo-iSPOR and Halo-iSPOR$^{Q251E}$ in WT and *ftsN*$^{\Delta SPOR+}$ strains. As a control, Halo-tag didn't localize to the septum. Scale bars, 5 µm. (**b**) Quantification of cell numbers with visible fluorescent band at the septum and the percentage of fluorescent intensity in septum and in the whole cell.

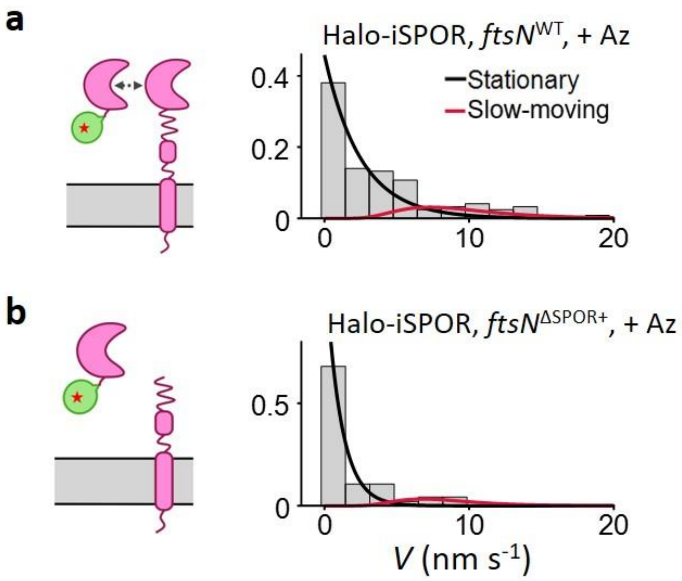


**Supplementary Fig. 5. Aztreonam treatment stopped the processive moving of Halo-iSPOR in both WT and *ftsN*$^{\Delta SPOR+}$ strains.**

Speed distribution of single Halo-iSPOR molecules in WT (**a**) and *ftsN*$^{\Delta SPOR+}$ (**b**) cells treated with aztreonam, overlaid with the fit curves of the stationary (black) and slow-moving (red) populations. Schematic representation of the fusion and strain background are shown as insets.

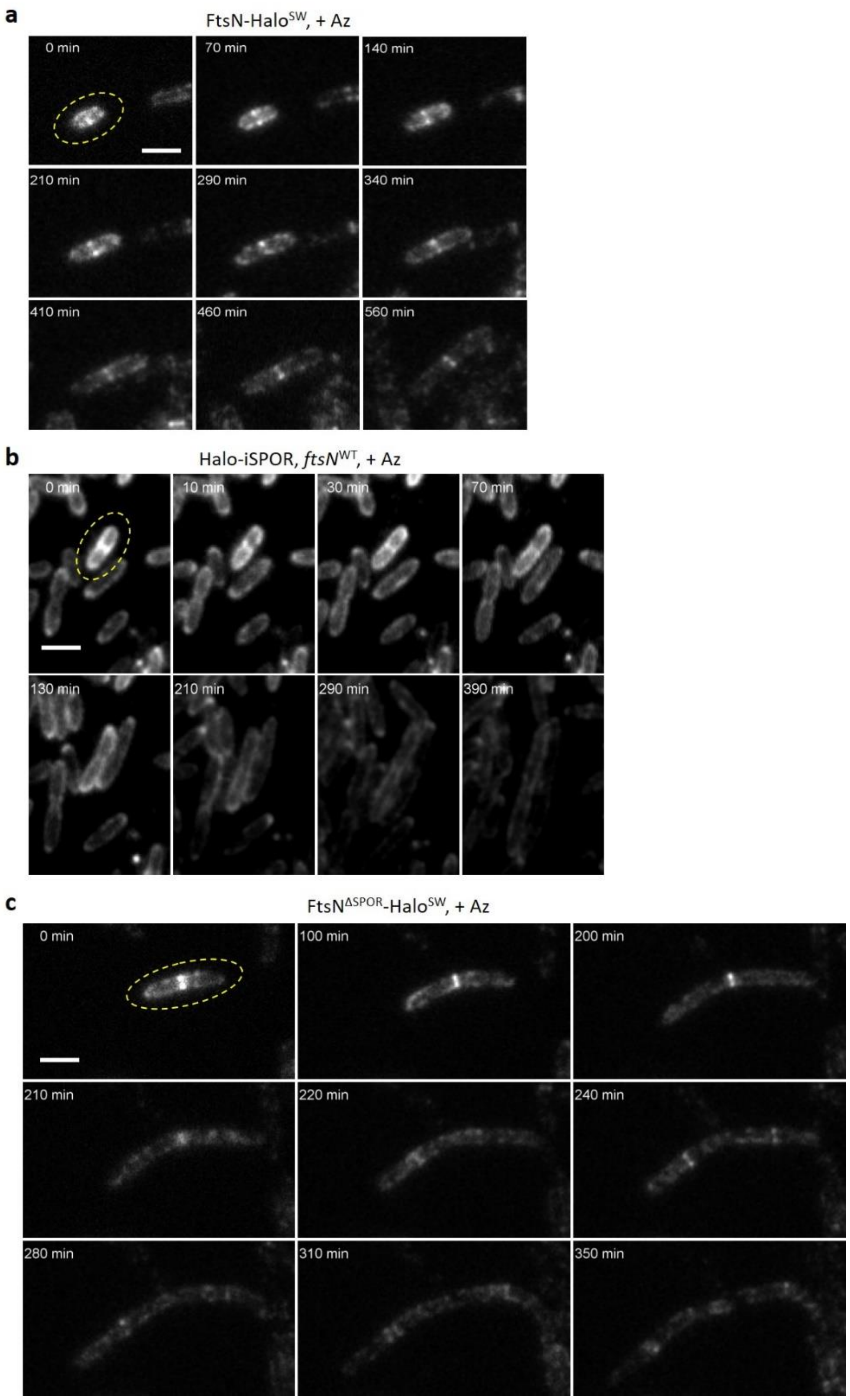


**Supplementary Fig. 6. Time-lapse images of cells treated with aztreonam.**

Time-lapse images of FtsN-Halo$^{SW}$ fusion cells (**a**), Halo-iSPOR fusion in WT cells (**b**), and FtsN$^{\Delta SPOR}$-Halo$^{SW}$ fusion cells (**c**) treated with aztreonam. The cells were highlighted with yellow dash oval. Scale bars, 3 µm. Only FtsN-Halo$^{SW}$ stayed at the septum while Halo-iSPOR and FtsN$^{\Delta SPOR}$-Halo$^{SW}$ left the septum when cells were treated with aztreonam.

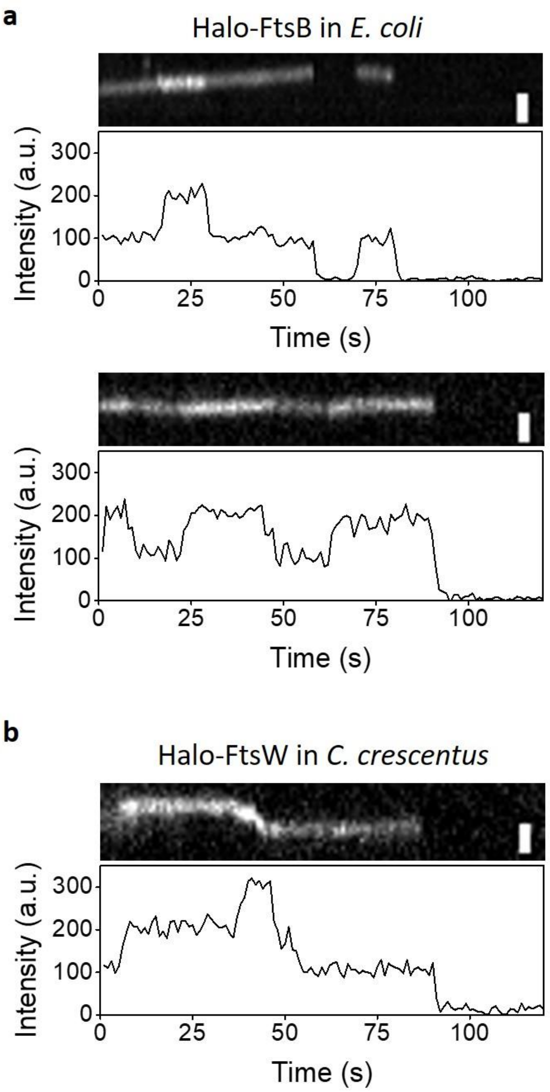


**Supplementary Fig. 7. FtsB and FtsW multimers observed in the septum of *E. coli* and *C. crescentus* cells.**

(**a**) Representative kymographs (top) and intensity traces (bottom) of two Halo-FtsB multimers in the septum of *E. coli.* Scale bars, 500 nm, (**b**) Representative kymographs (top) and intensity traces (bottom) of one Halo-FtsW multimer in the septum of *C. crescentus*. Scale bar, 300 nm.

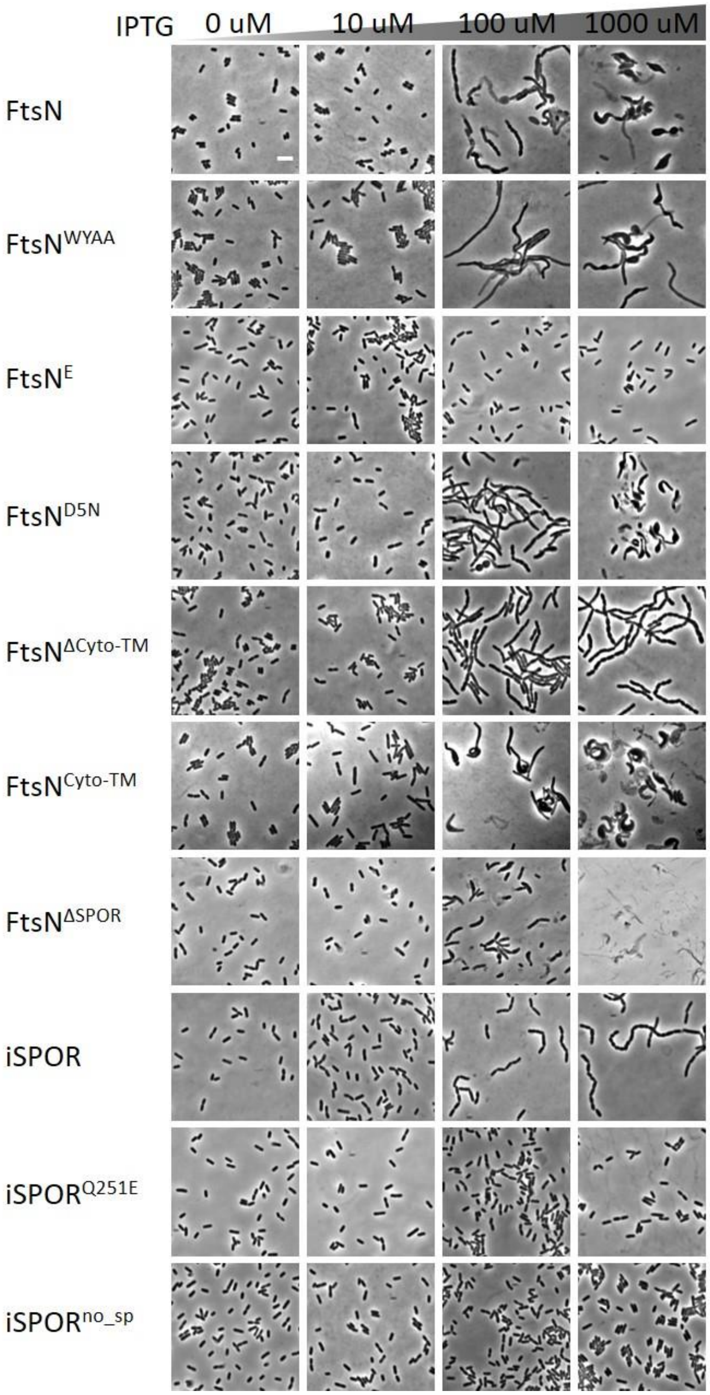


**Supplementary Fig. 8. Overexpression of iSPOR caused the chaining cell phenotype.**

Cell morphology of WT cells harbored different FtsN mutant plasmids induced at 0, 10, 100, and 1000 µM IPTG concentrations. no_sp means no signal peptide. Scale bar, 5 µm.

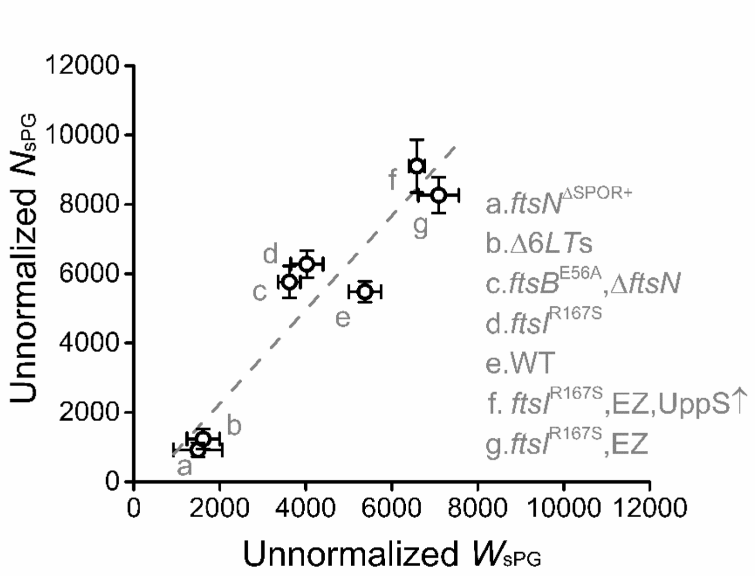


**Supplementary Fig. 9. The occupancy of FtsN-Halo$^{SW}$ ($N_{sPG}$ in *y* axis) and that of FtsW-RFP ($W_{sPG}$ in *x* axis) in a variety of strain backgrounds are highly correlated. Data were from this work and references[4, 6].**

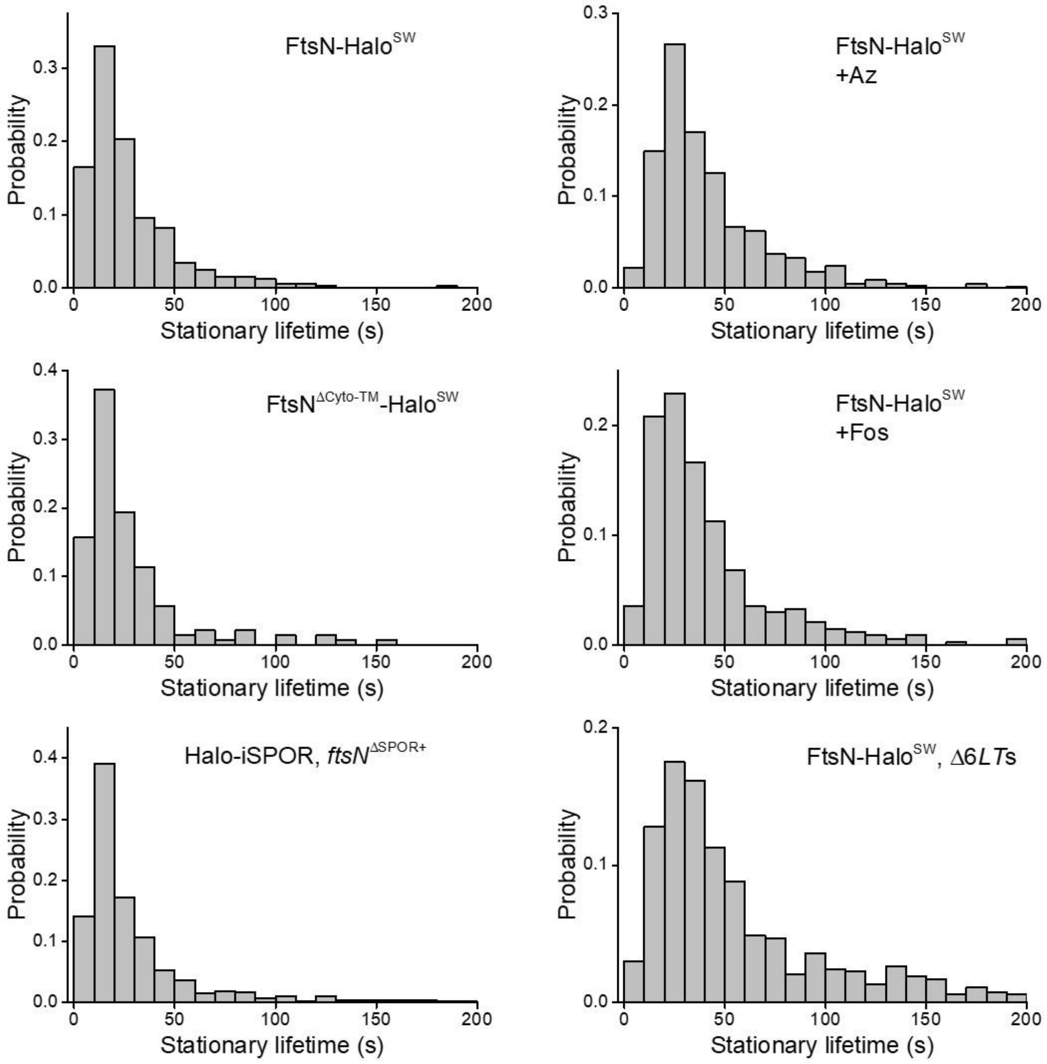


**Supplementary Fig. 10. The stationary lifetime distributions of FtsN variants under different conditions.**

The stationary lifetime distributions of FtsN variants under various conditions exhibited non-exponential distributions.

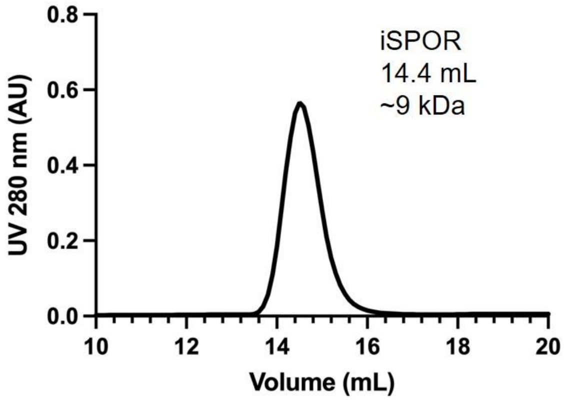


**Supplementary Fig. 11. Size exclusion chromatography showed that iSPOR majorly exists as a monomer in vitro.**

The molecular weight of monomeric iSPOR predicted from sequence is 10.5 kDa.

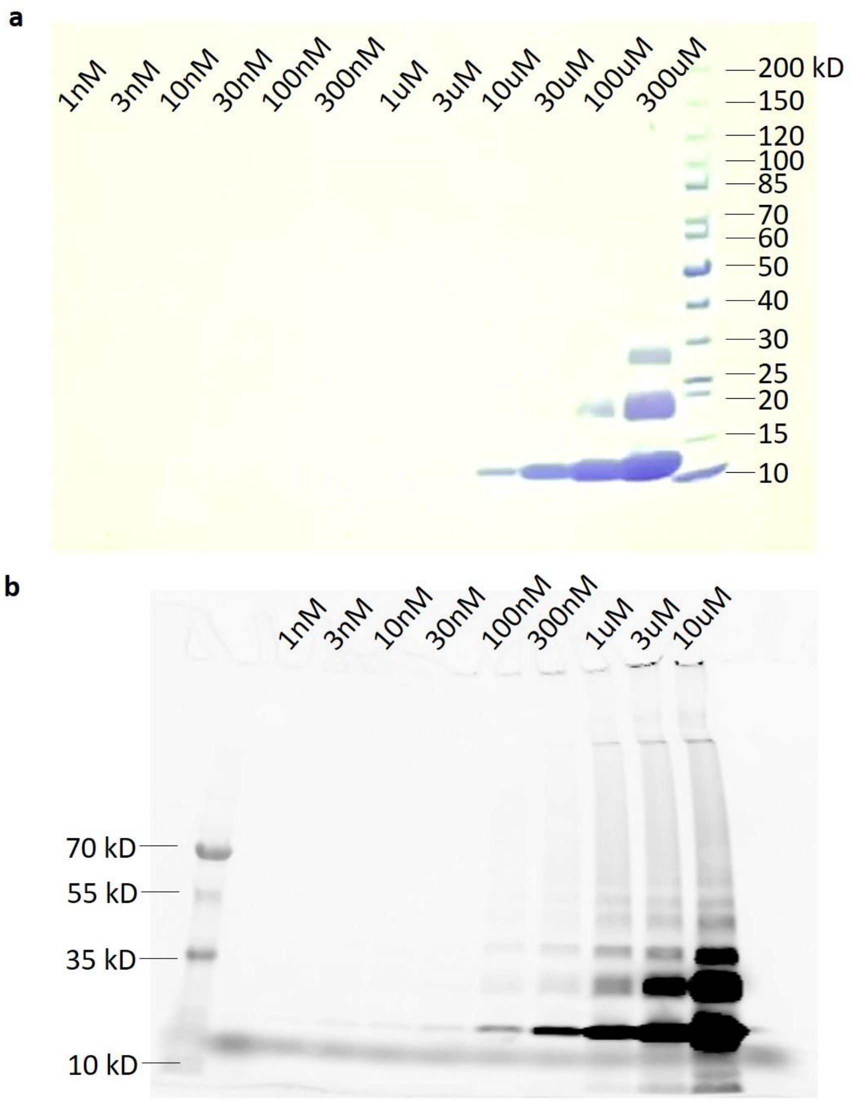


**Supplementary Fig. 12. iSPOR formed higher order oligomers at high concentrations.**

(**a**) Different concentrations of purified iSPOR protein were cross-linked with 0.1% glutaraldehyde for 15 min at RT. The protein gel was stained with Coomassie R-250. (**b**) Different concentrations of Cy3B-labeled purified iSPOR protein (iSPOR-Cy3B) were crosslinked with 0.1% glutaraldehyde for 15 min at RT. The protein gel was imaged on a Typhoon biomolecular imager directly without Coomassie staining.

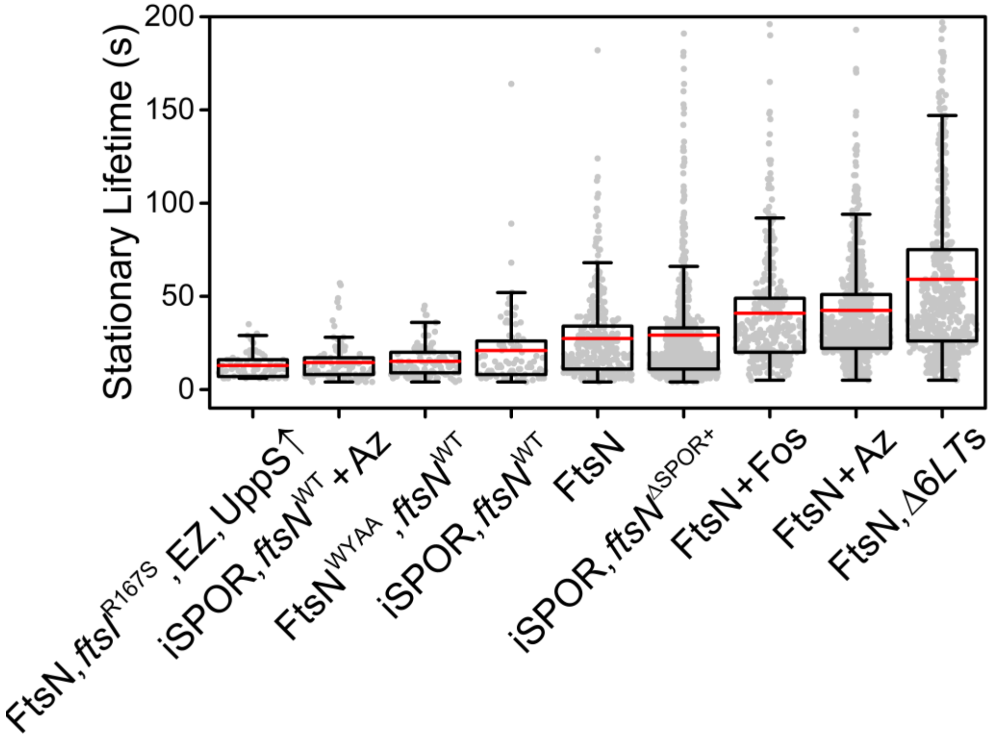


**Supplementary Fig. 13. Stationary lifetime of FtsN and variants under conditions where the expected levels of dnG for SPOR-binding varied.**

**Supplementary Table 1. Strains used in this study.**

| **Strains** | **Relevant genetic markers or features** | **Source, reference or construction** |
|---|---|---|
| **Stellar** | *mcrA* Δ(*mrr-hsdRMS-mcrBC*) φ80(*lacZ*)ΔM15 Δ(*lacZYA-argF*)*U169 endA1 recA1 supE44 thi-1 gyrA96 relA1* | Takara, cloning host |
| **MG1655** | *ilvG rfb*50 *rph1* | [13] |
| ***E. coli* Shuffle T7** | F´ *lac, pro,* $lacI^q$ */ Δ(ara-leu)7697 araD139 fhuA2 lacZ::T7 gene1 Δ(phoA)PvuII phoR ahpC* galE (or U) galK λatt::pNEB3-r1-cDsbC* (Spec$^R$, $lacI^q$) *ΔtrxB rpsL150*(Str$^R$) *Δgor Δ(malF)3* | New England Biolabs, expression host |
| **BW25113** | *rrnB3* Δ*lacZ4787 hsdR514* Δ(*araBAD*)567 Δ(*rhaBAD*)*568 rph-1* | [14] |
| **OmniMAX-2 T1R** | F' [$proAB^+$ $lacI^q$ *lacZ*ΔM15 Tn*10*(Tet$^r$) Δ(*ccdAB*)] *mcrA* Δ(*mrr-hsdRMS-mcrBC*) φ80(*lacZ*)ΔM15 Δ(*lacZYA-argF*) *U169 endA1 recA1 supE44 thi-1 gyrA96 relA1 tonA panD* | Invitrogen, cloning host |
| **EC251** | WT, Weiss lab isolate of MG1655 | [15] |
| **EC1908** | EC251 $P_{BAD}$::*ftsN* (Kan$^r$) | [16] |
| **EC2969** | BW25113 $ftsN^{\Delta SPOR}$<>*kan* | Recombineering |
| **EC2971** | EC251 $ftsN^{\Delta SPOR}$<>*kan* | P1 EC2969 X EC251, select Kan$^r$ |
| **EC3185** | EC251 $ftsN^{\Delta SPOR}$<>*frt* | pCP20 used to evict Kan$^r$ from EC2971 |
| **EC3338** | EC251 $ftsN^{\Delta SPOR}$<>*frt* | Independent isolate of E3185 |
| **EC3486** | EC251 Δ*amiA*<>*frt* Δ*amiC*<>*frt* | [17] |
| **TB170** | MG1655 Δ*amiB*<>*kan* | [18] |
| **EC3708** | EC251 Δ*rlpA*<>*frt* Δ*mltA*<>*frt* Δ*slt*<>*frt* Δ*mltD*<>*frt* Δ*mltE*<>*frt* Δ*mltC*<>*frt* | [17] |
| **EC5230** | EC251 $attP_{φ80}$::pDSW2083 [$P_{204\_7A}$::$ftsN\text{-}Halo^{SW}$ Spc$^r$] | [4] |

| | | |
|---|---|---|
| **EC5234** | EC251 P$_{BAD}$::*ftsN* (Kan$^{r}$) *attP*$_{φ80}$::pDSW2083 [P$_{204_7A}$::*ftsN-Halo*$^{SW}$ Spc$^{r}$] | [4] |
| **EC5257** | EC251 P$_{BAD}$::*ftsN* (Kan$^{r}$) *attP*$_{φ80}$::pDSW2093 [P$_{204_7A}$::*ftsN*$^{ΔSPOR}$-*Halo*$^{SW}$ Spc$^{r}$] | Integrate pDSW2093 into EC1908 using pAH123 |
| **EC5259** | EC251 P$_{BAD}$::*ftsN* (Kan$^{r}$) *attP*$_{φ80}$::pDSW2095 [P$_{204_7A}$::*ftsN*$^{WYAA}$-*Halo*$^{SW}$ Spcr] | Integrate pDSW2095 into EC1908 using pAH123 |
| **EC5275** | EC251 P$_{BAD}$::*ftsN* (Kan$^{r}$) *attP*$_{φ80}$::pDSW2103 [P$_{204_7A}$::*dsbA*$^{ss}$-*Halo-FtsN*$^{SPOR}$ Spc$^{r}$] | Integrate pDSW2103 into EC1908 using pAH123 |
| **EC5291** | EC251 Δ*amiA<>frt* Δ*amiC<>frt* *attP*$_{φ80}$::pDSW2083 [P$_{204_7A}$::*ftsN-Halo*$^{SW}$ Spc$^{r}$] | P1 EC5230 X EC3486, select Spc$^{r}$ |
| **EC5345** | EC251 Δ*amiA<>frt* Δ*amiC<>frt* Δ*amiB<>kan* *attP*$_{Φ80}$:: pDSW2083 [P$_{204_7A}$::*ftsN-Halo*$^{SW}$ Spc$^{r}$] | P1 TB170 X EC5291, select Spc$^{r}$ |
| **EC5402** | EC251 *ftsN*$^{ΔSPOR}$ *attP*$_{φ80}$::pDSW2103 [P$_{204_7A}$::*dsbA*$^{ss}$-*Halo-ftsN*$^{SPOR}$ Spc$^{r}$] | Integrate pDSW2103 into EC3185 using pAH123 |
| **EC5415** | EC5402 / pDSW2119 [P$_{ftsN}$::*ftsN*$^{ΔSPOR}$ *ori*SC101 Kan$^{r}$] | Transformation |
| **JL248** | EC1908 / pJL108 [P$_{T5\text{-}lac}$::*ftsN-mNG*$^{SW}$ Amp$^{r}$] | [4] |
| **JL337** | EC251 *ftsN*$^{ΔSPOR<>frt}$ / pDSW2119 [P$_{ftsN}$::*ftsN*$^{ΔSPOR}$ *ori*SC101 Kan$^{r}$] | Transform pDSW2119 into EC3338 |
| **JL351** | EC251 / pJL119 [P$_{T5\text{-}lac}$::*ftsN* Cam$^{r}$] | Transformation |
| **JL352** | EC251 / pJL120 [P$_{T5\text{-}lac}$::*ftsN*$^{WYAA}$ Cam$^{r}$] | Transformation |
| **JL353** | EC251 / pJL121 [P$_{T5\text{-}lac}$::*ftsN*$^{ΔSPOR}$ Cam$^{r}$] | Transformation |
| **JL354** | EC251 / pJL122 [P$_{T5\text{-}lac}$::*ftsN*$^{Cyto\text{-}TM}$ Cam$^{r}$] | Transformation |
| **JL359** | EC251 / pJL123 [P$_{T5\text{-}lac}$::*dsbA*$^{ss}$-*ftsN*$^{SPOR}$ Cam$^{r}$] | Transformation |

| | | |
|---|---|---|
| **JL360** | EC251 / pJL124 [$P_{T5\text{-}lac}$::*dsbA*$^{ss}$-*ftsN*$^{SPOR\ ,Q251E}$ Cam$^{r}$] | Transformation |
| **JL362** | JL337 / pXY349 [*bla lacI*$^{Q1}$ $P_{T5\text{-}lac}$::*ftsW-tagRFP-t*] | Transformation |
| **JL370** | EC251 / pJL128 [$P_{T5\text{-}lac}$::*dsbA*$^{ss}$-*ftsN*$^{E}$ Cam$^{r}$] | Transformation |
| **JL378** | EC1908 / pJL130 [$P_{T5\text{-}lac}$::*ftsN*$^{Q251E}$-*Halo*$^{SW}$ Amp$^{r}$] | Transformation |
| **JL386** | EC251 / pJL131 [$P_{T5\text{-}lac}$::*ftsN*$^{Cyto\text{-}TM,\ D5N}$ Cam$^{r}$] | Transformation |
| **JL431** | EC251 / pJL139 [$P_{T5\text{-}lac}$::*dsbA*$^{ss}$-*Halo-ftsN*$^{SPOR,Q251E}$ Cam$^{r}$] | Transformation |
| **JL432** | EC5415 / pJL137 [$P_{T5\text{-}lac}$::*dsbA*$^{ss}$-*Halo-ftsN*$^{SPOR}$ Cam$^{r}$] | This study |
| **JL435** | EC3708 / pJL132 [$P_{T5\text{-}lac}$::*ftsN-Halo*$^{SW}$ Cam$^{r}$] | This study |

**Supplementary Table 2: Plasmids used in this study.**

| Plasmid | Relevant features/description | Source or reference |
|---|---|---|
| **pDSW1333** | $P_{T5\text{-}lac}$::His$_6$-*ftsN*$^{SPOR}$ Amp$^r$ | [12] |
| **pDSW1920** | $P_{204\_7A}$::*Halo-ftsN lacI*$^Q$ *oriR*$_\gamma$ *attP*$_{\Phi80}$ Spc$^r$ | [4] |
| **pDSW2035** | $P_{204\_7A}$::*ftsN-Halo*$^{Q141SW}$ *lacI*$^{Q}$* *oriR*$_\gamma$ *attP*$_{\Phi80}$ Spc$^r$ | [4] |
| **pDSW2083** | $P_{204\_7A}$::*ftsN-Halo*$^{SW}$ *lacI*$^{Q}$* *oriR*$_\gamma$ *attP*$_{\Phi80}$ Spc$^r$ | [4] |
| **pDSW2093** | $P_{204\_7A}$::*ftsN*$^{\Delta SPOR}$-*Halo*$^{SW}$ *lacI*$^{Q}$* *oriR*$_\gamma$ *attP*$_{\Phi80}$ Spc$^r$ | This study |
| **pDSW2095** | $P_{204\_7A}$::*ftsN*$^{WYAA}$-*Halo*$^{SW}$ *lacI*$^{Q}$* *oriR*$_\gamma$ *attP*$_{\Phi80}$ Spc$^r$ | This study |
| **pDSW2099** | $P_{204\_7A}$::*dsbA*$^{ss}$-*Halo-ftsN*$^{\Delta Cyto\text{-}TM}$ *lacI*$^{Q}$* *oriR*$_\gamma$ *attP*$_{\Phi80}$ Spc$^r$ | [4] |
| **pDSW2119** | $P_{ftsN}$::*ftsN*$^{\Delta SPOR}$ *ori*SC101 Kan$^r$ | This study |
| **pJL019** | $P_{T5\text{-}lac}$::*mNG-ftsN*, Amp$^r$ | [4] |
| **pJL062** | $P_{204\_7A}$::*Halo-ftsN*$^{WYAA}$ *lacI*$^Q$ *oriR*$_\gamma$ *attP*$_{\Phi80}$ Spc$^r$ | This study |
| **pJL074** | $P_{204\_7A}$::*dsbA*$^{ss}$-*Halo-ftsN*$^{SPOR}$ *lacI*$^{Q}$* *oriR*$_\gamma$ *attP*$_{\Phi80}$ Spc$^r$ | [4] |
| **pJL077** | $P_{204\_7A}$::*mNG-ftsN*$^{WYAA}$ *lacI*$^Q$ *oriR*$_\gamma$ *attP*$_{\Phi80}$ Spc$^r$ | This study |
| **pJL108** | $P_{T5\text{-}lac}$::*mNG-ftsN*$^{SW}$ Amp$^r$ | [4] |
| **pJL113** | $P_{T5\text{-}lac}$::*ftsN-Halo*$^{SW}$, Amp$^r$ | [4] |
| **pJL119** | $P_{T5\text{-}lac}$::*ftsN* Cam$^r$ | [4] |
| **pJL120** | $P_{T5\text{-}lac}$::*ftsN*$^{WYAA}$ Cam$^r$ | This study |
| **pJL121** | $P_{T5\text{-}lac}$::*ftsN*$^{\Delta SPOR}$ Cam$^r$ | This study |
| **pJL122** | $P_{T5\text{-}lac}$::*ftsN*$^{Cyto\text{-}TM}$ Cam$^r$ | This study |
| **pJL123** | $P_{T5\text{-}lac}$::*dsbA*$^{ss}$-*ftsN*$^{SPOR}$ Cam$^r$ | This study |
| **pJL124** | $P_{T5\text{-}lac}$::*dsbA*$^{ss}$-*ftsN*$^{SPOR,Q251E}$ Cam$^r$ | This study |
| **pJL128** | $P_{T5\text{-}lac}$::*dsbA*$^{ss}$-*ftsN*$^{E}$ Cam$^r$ | This study |
| **pJL130** | $P_{T5\text{-}lac}$::*ftsN*$^{Q251E}$-*Halo*$^{SW}$ Amp$^r$ | This study |
| **pJL131** | $P_{T5\text{-}lac}$::*ftsN*$^{Cyto\text{-}TM,\ D5N}$ Cam$^r$ | This study |
| **pJL132** | $P_{T5\text{-}lac}$::*Halo-ftsN*$^{SW}$ Cam$^r$ | [4] |
| **pJL137** | $P_{T5\text{-}lac}$::*dsbA*$^{ss}$-*Halo-ftsN*$^{SPOR}$ Cam$^r$ | This study |
| **pJL139** | $P_{T5\text{-}lac}$::*dsbA*$^{ss}$-*Halo-ftsN*$^{SPOR,Q251E}$ Cam$^r$ | This study |
| **pXY349** | *bla lacI*$^{Q1}$ $P_{T5\text{-}lac}$::*ftsW-tagRFP-t* | [6] |

**Deletion of *ftsN*$^{SPOR}$ by recombineering.** FtsN's SPOR domain codons 252-313 were replaced by λRwith a kanamycin resistance cassette PCR-amplified from pKD13 using primers P1752 and P1753.

**Plasmid construction.**

**pDSW2093.** Primers P2433 and P2182 were used to PCR-amplify an *ftsN-Halo*$^{SW}$ fusion spanning codons 1-242 of *ftsN* followed by a TAA stop codon, thus deleting the SPOR domain, which starts at codon 240. The template was pDSW2083. The 1855 bp DNA fragment was digested with AflII and KpnI, then ligated into pDSW2035 that had been cut with the same enzymes.

**pDSW2095.** Primers P2461 and P2462 were used to PCR-amplify the *ftsN*(W83A Y85A) double substitution allele from pJL077. The 839 bp fragment was digested with AccI and KpnI, then ligated into pDSW2083 that had been cut with the same enzymes.

**pDSW2103.** Primers P2460 and P2467 were used to PCR-amplify the *ftsN* SPOR domain with pDSW2083 as template. The 383 bp fragment was digested with AccI and KpnI, then ligated into pDSW2099 that had been cut with the same enzymes.

**pDSW2119.** This plasmid is a pSC101 derivative that mildly overexpresses *ftsN*$^{\Delta SPOR}$ from its native promoter, $P_{ftsN}$. Insert was generated PCR-amplifying a 986 bp DNA fragment that extends from 200 bp upstream of *ftsN* through codon 242, followed by a TGA stop codon. The primers were P2486 and P2487. The template was EC251 genomic DNA. A 2810 bp vector fragment was generated by PCR amplification of pHGx using primers P2484 and P2485.The fragments were joined using NEBuilder HiFi DNA Assembly Master Mix.

**pJL062**. The pJL062 plasmid was constructed from the pDSW1920 plasmid using the QuikChange protocol (Agilent) with the primers 60 and 61 to mutate the nucleotide sequence encoding for W83A, Y85A.

**pJL077**. Amplify *ftsN*$^{WYAA}$ gene with the vector backbone from pJL062 with primers 13 and 99. Amplify *mNeonGreen* gene from pJL019 with primers 12 and 100. The two DNA fragments were then joined by the In-Fusion Cloning Kit to generate the plasmid pJL077.

**pJL120**. The pJL120 plasmid was constructed from the pJL119 plasmid using the QuikChange protocol (Agilent) with the primers 60 and 61 to mutate the nucleotide sequence encoding for W83A, Y85A.

**pJL121**. Amplify the vector backbone from pJL119 with primers 72 and 126. Amplify *ftsN*$^{\Delta SPOR}$ gene from pDSW1920 with primers 65 and 127. The two DNA fragments were then joined by the In-Fusion Cloning Kit to generate the plasmid pJL121.

**pJL122**. Amplify the vector backbone from pJL119 with primers 72 and 126. Amplify *ftsN*$^{Cyto\text{-}TM}$ gene from pDSW1920 with primers 117 and 127. The two DNA fragments were then joined by the In-Fusion Cloning Kit to generate the plasmid pJL122.

**pJL123**. Amplify *ftsN*$^{SPOR}$ gene with the vector backbone from pJL119 with primers 72 and 106. Amplify *dsbA*$^{ss}$ gene from pDSW2099 with primers 181 and 183. The two DNA fragments were then joined by the In-Fusion Cloning Kit to generate the plasmid pJL123.

**pJL124**. The pJL124 plasmid was constructed from the pJL123 plasmid using the QuikChange protocol (Agilent) with the primers 62 and 63 to mutate the nucleotide sequence encoding for Q251E.

**pJL128**. Amplify *dsbA*$^{ss}$ gene with the vector backbone from pJL123 with primers 116 and 126. Amplify *ftsN*$^{E}$ gene from pDSW1920 with primers 186 and 187. The two DNA fragments were then joined by the In-Fusion Cloning Kit to generate the plasmid pJL128.

**pJL130**. The pJL130 plasmid was constructed from the pJL113 plasmid using the QuikChange protocol (Agilent) with the primers 62 and 63 to mutate the nucleotide sequence encoding for Q251E.

**pJL131**. The pJL131 plasmid was constructed from the pJL122 plasmid using the QuikChange protocol (Agilent) with the primers 168 and 169 to mutate the nucleotide sequence encoding for D5N.

**pJL137**. Amplify the vector backbone from pJL119 with primers 72 and 126. Amplify *dsbA*$^{ss}$*-Halo-ftsN*$^{SPOR}$ gene from pJL074 with primers 146 and 181. The two DNA fragments were then joined by the In-Fusion Cloning Kit to generate the plasmid pJL137.

**pJL139**. The pJL139 plasmid was constructed from the pJL137 plasmid using the QuikChange protocol (Agilent) with the primers 62 and 63 to mutate the nucleotide sequence encoding for Q251E.

**Supplementary Table 3. Oligonucleotides used in this study.**

| Name | Sequence (5’->3’) | Comment |
|---|---|---|
| **P1752** | CGCCAAAACCGACGGCGGAGAAAAAAGACGAACGCCGCTGGATGGTGCAGATTCCGGGGATCCGTCGACC | Forward primer to delete FtsN SPOR domain on chromosome of EC251 |
| **P1753** | AATTATAGATGGGGGGGATTTTGAGGGTTTCAACCCCCGGCGGCGAGCCGTGTAGGCTGGAGCTGCTTCG | Reverse primer to delete FtsN SPOR domain on chromosome of EC251 |
| **P2182** | GGCTGTGCAGGTCGTAAATC | Forward primer to construct pDSW2093 |
| **P2433** | GTGGGTACCTTATTTCTCCGCCGTCGGTTTTGG | Reverse primer to construct pDSW2093 |
| **P2460** | GCCTACCCGGATATTATCGTG | Reverse primer to construct pDSW2103 |
| **P2461** | GGCTGTCTACTCTGGAGATTTCCGGTGGTGGCGGTGGTAGCGAGTCCGAGACGCTGCAA | Forward primer to construct pDSW2095 |
| **P2462** | CTCGGTACCCGGGGATCCTTAACCCCCGGCGGCGAGCCGAATGCAGTTTG | Reverse primer to construct pDSW2095 |
| **P2484** | AGCTTACTGGCCGTCGTTTTACAACGTCGTG | Forward primer to generate vector for constructing pDSW2119 |
| **P2485** | GAGCTAACTCACATTATGGACAG | Reverse primer to generate vector for constructing pDSW2119 |
| **P2486** | AATGTGAGTTAGCTCGATATCGGAAGCTATGCTGTTATTGCTTGATC | Forward primer to generate insert for constructing pDSW2119 |
| **P2487** | GACGGCCAGTAAGCTTCATTTCTCCGCCGTCGGT | Reverse primer to generate insert for constructing pDSW2119 |
| **12** | GCTTTTTTGTACAAACTTGTTGATTTTATACAGTTCATCCATGCCCATC | Reverse primer to clone *mNG* into pJL077 |

| | | |
|---|---|---|
| **13** | ATCAACAAGTTTGTACAAAAAAGCAGG | Forward primer to amplify *ftsN*-vector fragment for pJL077 |
| **60** | CCAGAAGAACGCGCTCGCGCCATTAAAGAGCTG | Forward primer for site-specific mutagenesis into WYAA for pJL120 |
| **61** | CAGCTCTTTAATGGCGCGAGCGCGTTCTTCTGG | Reverse primer for site-specific mutagenesis into WYAA for pJL120 |
| **62** | CGCTGGATGGTGGAGTGCGGTTCGTTCAG | Forward primer for site-specific mutagenesis into Q251E for pJL124, pJL130 |
| **63** | CTGAACGAACCGCACTCCACCATCCAGCG | Reverse primer for site-specific mutagenesis into Q251E for pJL124, pJL130 |
| **65** | CCCTTAGCGGCCGCTCATTTCTCCGCCGTCGGTTTTG | Reverse primer to clone *ftsN*$^{\Delta SPOR}$ into pJL121 |
| **72** | ACTAGTAGTTAATTTCTCCTCTTTAATG | Reverse primer to amplify vector fragment for pJL121, pJL122, pJL123 |
| **99** | GGTCTGTTTCCTGTGTCTTAAGAAT | Reverse primer to amplify *ftsN*-vector fragment for pJL077 |
| **100** | CACAGGAAACAGACCATGGTGAGCAAAGGCGAAGAAG | Forward primer to clone *mNG* into pJL077 |
| **106** | AAAGACGAACGCCGCTGGATGG | Forward primer to amplify vector fragment for pJL123 |
| **116** | CGCCGATGCGCTAAACGCT | Reverse primer to amplify vector fragment for pJL128 |
| **117** | CGACCCTTAGCGGCCGCTCACGTAATGAAGTACAGACCACC | Reverse primer to clone *ftsN*$^{Cyto\text{-}TM}$ into pJL122 |

| | | |
|---|---|---|
| **126** | GCGGCCGCTAAGGGTCG | Forward primer to amplify vector fragment for pJL121, pJL122, pJL128 |
| **127** | GAGGAGAAATTAACTACTAGTATGGCACAACGAGATTATGTACGC | Forward primer to clone *ftsN*$^{\Delta SPOR}$ into pJL121, pJL122 |
| **146** | GACCCTTAGCGGCCGCTCAACCCCCGGCGGCGAGC | Reverse primer to clone *dsbA*$^{ss}$-*Halo-ftsN*$^{SPOR}$ into pJL137 |
| **168** | ATGGCACAACGAAATTATGTACGCCGCAGCCAAC | Forward primer for site-specific mutagenesis into D5N for pJL131 |
| **169** | GTTGGCTGCGGCGTACATAATTTCGTTGTGCCAT | Reverse primer for site-specific mutagenesis into D5N for pJL131 |
| **181** | CATTAAAGAGGAGAAATTAACTACTAGTATGAAAAAGATTTGGCTGGCGCTG | Forward primer to clone *dsbA*$^{ss}$ into pJL123, pJL137 |
| **183** | CTGCACCATCCAGCGGCGTTCGTCTTTCGCCGATGCGCTAAACGCTAA | Reverse primer to clone *dsbA*$^{ss}$ into pJL123 |
| **186** | GTTTAGTTTTAGCGTTTAGCGCATCGGCGACCGGAAACGGACTACCACC | Forward primer to clone *ftsN*$^{E}$ into pJL128 |
| **187** | CTGCAGGTCGACCCTTAGCGGCCGCTTAACCGGCAGAAGGTTCTGTG | Reverse primer to clone *ftsN*$^{E}$ into pJL128 |

**Supplementary Table 4. Dynamics of FtsN and FtsW in WT, Δ*amiABC*, and Δ6*LT*s strains.**

| Genotype | Name | $P_{stat}$[c] (%) ($n_{stat}$) | $P_{sm}$[c] (%) ($n_{sm}$) | $P_{fm}$[c] (%) ($n_{fm}$) | $V_{sm}$[d] (nm $s^{-1}$) | $V_{fm}$[d] (nm$s^{-1}$) | $T_{stat}$[d] (s) | $T_{sm}$[d] (s) | $T_{fm}$[d] (s) | $PL_{sm}$[d] (nm) | $PL_{fm}$[d] (nm) |
|---|---|---|---|---|---|---|---|---|---|---|---|
| MG1655[b] | FtsN-Halo$^{SW}$ | 55.1 ± 2.2 (315) | 44.9 ± 1.6 (256) | 0 | 9.4 ± 0.2 | N.A.[a] | 27.3 ± 1.3 | 14.5 ± 0.7 | N.A.[a] | 122 ± 5 | N.A.[a] |
| | FtsW-RFP | 54.9 ± 1.7 (258) | 33.4 ± 2.3 (157) | 11.7 ± 1.8 (55) | 9.9 ± 0.1 | 43.7 ± 0.1 | 18.2 ± 0.8 | 20.1 ± 0.1 | 8.3 ± 0.3 | 161 ± 2 | 287 ± 5 |
| MG1655, Δ*amiABC* | FtsN-Halo$^{SW}$ | 58.6 ± 2.3 (109) | 0 | 41.4 ± 1.9 (77) | N.A.[a] | 37.9 ± 3.4 | 9.4 ± 0.6 | N.A.[a] | 6.4 ± 0.5 | N.A.[a] | 195 ± 14 |
| | FtsW-RFP | 61.7 ± 3.5 (116) | 0 | 38.3 ± 2.7 (72) | N.A.[a] | 39.0 ± 3.6 | 9.1 ± 0.6 | N.A.[a] | 6.7 ± 0.5 | N.A.[a] | 205 ± 15 |
| MG1655, Δ6*LT*s | FtsN-Halo$^{SW}$ | 91.2 ± 2.6 (546) | 8.8 ± 2.0 (53) | 0 | 8.4 ± 0.4 | N.A.[a] | 59.1 ± 2.1 | 18.1 ± 1.5 | N.A.[a] | 140 ± 10 | N.A.[a] |
| | FtsW-RFP | 88.4 ± 2.7 (252) | 11.6 ± 2.5 (33) | 0 | 8.3 ± 0.5 | N.A.[a] | 23.9 ± 1.1 | 17.4 ± 1.5 | N.A.[a] | 139 ± 13 | N.A.[a] |

All the experiments were performed with cells grown in M9-glucose minimal medium.

[a] N.A. means not applicable.

[b] In the MG1655 WT background, the SMT data for FtsN-Halo$^{SW}$ was from reference[4], the SMT data for FtsW-RFP was from reference[6] and was reanalyzed using speed deconvolution.

[c] Percentage of the segment number of FtsN or FtsW molecules grouped in stationary state ($P_{stat}$), slow-moving state ($P_{sm}$), and fast-moving state ($P_{fm}$). $P_{sm}$ and $P_{fm}$ were obtained from peed deconvolution. Data are presented as mean ± error, where the error is the standard deviation from 500 bootstrap samples pooled from three independent experiments. *n* is the segment number in each state.

[d] Average speed ($V_{sm}$ *and* $V_{fm}$), stationary lifetime ($T_{stat}$), persistent running time ($T_{sm}$ and $T_{fm}$), and persistent running length ($PL_{sm}$ *and* $PL_{fm}$) of FtsN or FtsW molecules in different states. Data are presented as mean ± *s.e.m.*

**Supplementary Table 5. Dynamics of FtsN variants and FtsW in the strain background of truncation or mutation of FtsN's SPOR domain.**

| Genotype | Name | Drug or medium | $P_{stat}$[b] (%) ($n_{stat}$) | $P_{sm}$[b] (%) ($n_{sm}$) | $P_{fm}$[b] (%) ($n_{fm}$) | $V_{sm}$[c] (nm $s^{-1}$) | $V_{fm}$[c] ($nms^{-1}$) | $T_{stat}$[c] (s) | $T_{sm}$[c] (s) | $T_{fm}$[c] (s) | $PL_{sm}$[c] (nm) | $PL_{fm}$[c] (nm) |
|---|---|---|---|---|---|---|---|---|---|---|---|---|
| MG1655, *ftsN*$^{\Delta SPOR+}$ | FtsW-RFP | M9 | 73.5 ± 2.1 (353) | 7.9 ± 3.0 (38) | 18.5 ± 1.9 (89) | 9.7 ± 0.1 | 32.4 ± 0.1 | 10.4 ± 0.4 | 21.1 ± 0.5 | 9.4 ± 0.2 | 189 ± 5 | 209 ± 2 |
| | FtsN$^{\Delta SPOR}$-Halo$^{SW}$ | M9 | 40.3 ± 1.8 (85) | 13.3 ± 2.9 (28) | 46.4 ± 1.6 (98) | 9.4 ± 0.1 | 32.7 ± 0.1 | 9.2 ± 0.6 | 7.5 ± 0.1 | 5.5 ± 0.1 | 69 ± 2 | 173 ± 1 |
| | | Az | 43.3 ± 2.9 (139) | 0 | 56.7 ± 2.1 (182) | N.A.[a] | 22.9 ± 1.2 | 22.5 ± 1.5 | N.A.[a] | 9.7 ± 0.6 | N.A.[a] | 165 ± 6 |
| MG1655, $P_{BAD}$::*ftsN* | FtsN$^{Q251E}$-Halo$^{SW}$ | M9 | 41.8 ± 1.5 (81) | 25.3 ± 2.2 (49) | 33.0 ± 1.9 (64) | 10.3 ± 0.1 | 28.4 ± 0.1 | 13.1 ± 0.9 | 9.7 ± 0.2 | 6.4 ± 0.2 | 94 ± 3 | 164 ± 2 |

[a] N.A. means not applicable.

[b] Percentage of the segment number of FtsN variant or FtsW molecules grouped in stationary state ($P_{stat}$), slow-moving state ($P_{sm}$), and fast-moving state ($P_{fm}$). $P_{sm}$ and $P_{fm}$ were obtained from peed deconvolution. Data are presented as mean ± error, where the error is the standard deviation from 500 bootstrap samples pooled from three independent experiments. $n$ is the segment number in each state.

[c] Average speed ($V_{sm}$ *and* $V_{fm}$), stationary lifetime ($T_{stat}$), persistent running time ($T_{sm}$ and $T_{fm}$), and persistent running length ($PL_{sm}$ *and* $PL_{fm}$) of FtsN variant or FtsW molecules in different states. Data are presented as mean ± *s.e.m.*

**Supplementary Table 6. Dynamics of FtsN$^{WYAA}$ and iSPOR in WT or *ftsN*$^{\Delta SPOR+}$ strain backgrounds.**

| Genotype | Name | Drug or medium | $P_{stat}$[a] (%) ($n_{stat}$) | $P_{sm}$[a] (%) ($n_{sm}$) | $V_{sm}$[b] (nm $s^{-1}$) | $T_{stat}$[b] (s) | $T_{sm}$[b] (s) | $PL_{sm}$[b] (nm) |
|---|---|---|---|---|---|---|---|---|
| MG1655, $P_{BAD}$::*ftsN* | FtsN$^{WYAA}$-Halo$^{SW}$ | M9 | 56.4 ± 2.8 (102) | 43.6 ± 3.5 (79) | 9.6 ± 0.5 | 15.0 ± 0.9 | 10.7 ± 0.8 | 92 ± 5 |
| | Halo-iSPOR | M9 | 58.5 ± 2.1 (86) | 41.5 ± 2.9 (61) | 9.4 ± 0.5 | 20.9 ± 2.4 | 9.8 ± 0.7 | 84 ± 5 |
| | Halo-iSPOR | Az | 86.0 ± 2.1 (104) | 14.0 ± 3.8 (17) | 8.2 ± 0.8 | 14.5 ± 1.0 | 13.4 ± 2.0 | 95 ± 13 |
| | Halo-iSPOR$^{Q251E}$ | M9 | 63.7 ± 4.5 (58) | 36.3 ± 3.8 (33) | 10.8 ± 1.1 | 19.7 ± 1.2 | 9.6 ± 0.6 | 99 ± 9 |
| MG1655, *ftsN*$^{\Delta SPOR+}$ | Halo-iSPOR | M9 | 92.2 ± 1.3 (602) | 7.8 ± 2.1 (51) | 9.5 ± 0.5 | 29.2 ± 1.3 | 12.6 ± 0.9 | 111 ± 7 |
| | Halo-iSPOR | Az | 93.6 ± 2.5 (44) | 6.4 ± 2.8 (3) | 8.2 ± 1.0 | 32.6 ± 4.5 | 10.0 ± 0.6 | 81 ± 7 |

[a] Percentage of the segment number of FtsN$^{WYAA}$ or iSPOR molecules grouped in stationary state ($P_{stat}$) and slow-moving state ($P_{sm}$). Data are presented as mean ± error, where the error is the standard deviation from 500 bootstrap samples pooled from three independent experiments. *n* is the segment number in each state.

[b] Average speed ($V_{sm}$), stationary lifetime ($T_{stat}$), persistent running time ($T_{sm}$), and persistent running length ($PL_{sm}$) of FtsN$^{WYAA}$ or iSPOR molecules in different states. Data are presented as mean ± *s.e.m.*

**Supplementary Table 7. Static Light Scattering.**

| Sample | Radius (nm) | Polydispersity (PD) (%) | MW (kDa) |
|---|---|---|---|
| **FtsN$^{SPOR}$** | 2.0 | 14.8 | 12.00 ± 0.05 |

### Captions for Movies

**Supplementary Movie 1. Time-lapse movie of FtsN-Halo$^{SW}$ fusion cells treated with aztreonam.**
Images were acquired every 10 minutes for 560 minutes, with a 100-ms exposure time. Scale bar, 500 nm.

**Supplementary Movie 2. Time-lapse movie of Halo-iSPOR fusion cells treated with aztreonam.**
Images were acquired every 10 minutes for 210 minutes, with a 100-ms exposure time. Scale bar, 500 nm.

**Supplementary Movie 3. Time-lapse movie of FtsN$^{\Delta SPOR}$-Halo$^{SW}$ fusion cells treated with aztreonam.**
Images were acquired every 10 minutes for 360 minutes, with a 100-ms exposure time. Scale bar, 500 nm.

**Supplementary Movie 4. SMT image of a cell with a processively moving FtsN-Halo$^{SW}$ multimer.**
Images were acquired every 1 s for 220 s, with a 100-ms exposure time. Scale bar, 500 nm.

**Supplementary Movie 5. SMT image of a cell with a stationary FtsN-Halo$^{SW}$ multimer.**
Images were acquired every 1 s for 230 s, with a 100-ms exposure time. Scale bar, 500 nm.

**Supplementary Movie 6. SMT image of a cell with a processively moving Halo-FtsB multimer in *E. coli*.**
Images were acquired every 1 s for 120 s, with a 500-ms exposure time. Scale bar, 500 nm.

**Supplementary Movie 7. SMT image of a cell with a Halo-FtsB multimer exhibiting both stationary and processively moving states in *E. coli*.**

Images were acquired every 1 s for 120 s, with a 500-ms exposure time. Scale bar, 500 nm.

**Supplementary Movie 8. SMT image of a cell with a processively moving Halo-FtsW multimer in *C. crescentus*.**

Images were acquired every 1 s for 120 s, with a 500-ms exposure time. Scale bar, 500 nm.

### Supplementary References


1. Laboratory, C.S.H. M9 minimal medium (standard). *Cold Spring Harb Protoc* (2010).
2. Guzman, L.M., Belin, D., Carson, M.J. & Beckwith, J. Tight regulation, modulation, and high-level expression by vectors containing the arabinose $P_{BAD}$ promoter. *J. Bacteriol.* **177**, 4121-4130 (1995).
3. Haldimann, A. & Wanner, B.L. Conditional-replication, integration, excision, and retrieval plasmid-host systems for gene structure-function studies of bacteria. *J. Bacteriol.* **183**, 6384-6393 (2001).
4. Lyu, Z. *et al.* FtsN maintains active septal cell wall synthesis by forming a processive complex with the septum-specific peptidoglycan synthases in *E. coli*. *Nat. Commun.* **13**, 5751 (2022).
5. Lyu, Z., Coltharp, C., Yang, X. & Xiao, J. Influence of FtsZ GTPase activity and concentration on nanoscale Z-ring structure in vivo revealed by three-dimensional Superresolution imaging. *Biopolymers* **105**, 725-734 (2016).
6. Yang, X. *et al.* A two-track model for the spatiotemporal coordination of bacterial septal cell wall synthesis revealed by single-molecule imaging of FtsW. *Nat. Microbiol.* **6**, 584-593 (2021).
7. Ovesny, M., Krizek, P., Borkovec, J., Svindrych, Z. & Hagen, G.M. ThunderSTORM: a comprehensive ImageJ plug-in for PALM and STORM data analysis and super-resolution imaging. *Bioinformatics* **30**, 2389-2390 (2014).
8. Schneider, C.A., Rasband, W.S. & Eliceiri, K.W. NIH Image to ImageJ: 25 years of image analysis. *Nat. Methods* **9**, 671-675 (2012).
9. Huang, B., Wang, W., Bates, M. & Zhuang, X. Three-dimensional super-resolution imaging by stochastic optical reconstruction microscopy. *Science* **319**, 810-813 (2008).
10. Sbalzarini, I.F. & Koumoutsakos, P. Feature point tracking and trajectory analysis for video imaging in cell biology. *J. Struct. Biol.* **151**, 182-195 (2005).
11. McCausland, J.W. *et al.* Treadmilling FtsZ polymers drive the directional movement of sPG-synthesis enzymes via a Brownian ratchet mechanism. *Nat. Commun.* **12**, 609 (2021).
12. Duncan, T.R., Yahashiri, A., Arends, S.J., Popham, D.L. & Weiss, D.S. Identification of SPOR domain amino acids important for septal localization, peptidoglycan binding, and a disulfide bond in the cell division protein FtsN. *J. Bacteriol.* **195**, 5308-5315 (2013).
13. Guyer, M.S., Reed, R.R., Steitz, J.A. & Low, K.B. Identification of a sex-factor-affinity site in *E. coli* as gamma delta. *Cold Spring Harb Symp Quant Biol* **45 Pt 1**, 135-140 (1981).
14. Baba, T. *et al.* Construction of *Escherichia coli* K-12 in-frame, single-gene knockout mutants: the Keio collection. *Mol. Syst. Biol.* **2**, 0008 (2006).
15. Arends, S.J. & Weiss, D.S. Inhibiting cell division in *Escherichia coli* has little if any effect on gene expression. *J. Bacteriol.* **186**, 880-884 (2004).
16. Tarry, M. *et al.* The Escherichia coli cell division protein and model Tat substrate SufI (FtsP) localizes to the septal ring and has a multicopper oxidase-like structure. *J. Mol. Biol.* **386**, 504-519 (2009).
17. Yahashiri, A., Jorgenson, M.A. & Weiss, D.S. Bacterial SPOR domains are recruited to septal peptidoglycan by binding to glycan strands that lack stem peptides. *Proc. Natl Acad. Sci. USA* **112**, 11347-11352 (2015).

18. Uehara, T., Parzych, K.R., Dinh, T. & Bernhardt, T.G. Daughter cell separation is controlled by cytokinetic ring-activated cell wall hydrolysis. *EMBO J.* **29**, 1412-1422 (2010).
